# Muc5b-deficient mice develop early histological lung abnormalities

**DOI:** 10.1101/705962

**Authors:** Hélène Valque, Valérie Gouyer, Catherine Duez, Christophe Leboeuf, Philippe Marquillies, Marc Lebert, Ségolène Plet, Bernhard Ryffel, Anne Janin, Frédéric Gottrand, Jean-Luc Desseyn

## Abstract

Gel-forming mucins are the main organic component responsible for physical properties of the mucus hydrogels. While numerous biological functions of these mucins are well documented, specific physiological functions of each mucin are largely unknown. To investigate *in vivo* functions of the gel-forming mucin Muc5b, which is with Muc5ac the major secreted airway mucin, we generated mice in which Muc5b was disrupted and maintained in the absence of environmental stress. Adult Muc5b-deficient mice displayed bronchial hyperplasia and metaplasia, interstitial thickening, alveolar collapse, immune cell infiltrates, fragmented and disorganized elastin fibers and collagen deposits that were for approximately one fifth of mice associated with altered pulmonary function leading to respiratory failure. These lung abnormalities start early in life, as demonstrated for one fourth 2-day-old Muc5b-deficient pups. Thus, the mouse mucin Muc5b is essential for maintaining normal lung function.

## INTRODUCTION

Mucus gel is the first line of defense in the lung. Gel-forming mucins are high molecular weight macromolecules that make up the major organic component of mucus. These molecules are heavily *O*-glycosylated and dimerize through their carboxy-terminal region and polymerize through their amino-terminal region. Long polymers are secreted at the cell surface and form mucus gel when in contact with water (Thornton *et al.*, 2008). The central region of gel-forming mucins is enriched with proline and hydroxyl amino acids (Ser/Thr/Pro regions) and is extensively *O*-glycosylated. Five gel-forming mucins, MUC2, -6, -5AC, -5B and -19, have been cloned and characterized and are highly conserved between humans and rodents. They all contain a large exon encoding the Ser/Thr/Pro region.

In lung, the mucus layer covers and protects the cell surface of the airways epithelium and traps exogenous particles and microorganisms for mucociliary clearance. In humans, MUC5AC is secreted by goblet cells whereas MUC5B is secreted by submucosal glands. MUC5B is also secreted by the salivary glands, nasal mucosa, lacrimal glands, gallbladder, middle ear, submucosal glands of the trachea and esophagus, and the epithelium and glands of the endocervix. A similar pattern of expression was found in mouse tissues (Valque *et al.*, 2011; Portal *et al.*, 2017*b*) with the early expression of *MUC5B* during human development (Buisine *et al.*, 1999, 2000) and in mouse lungs at embryonic (E) day 12.5 or earlier (Portal *et al.*, 2017*b*).

The redundancy of the two gel-forming mucins in the lung make difficult to understand the precise function of each mucin. Dysregulation of *MUC5B* expression has been reported in airway diseases (Rose & Voynow, 2006; Fahy & Dickey, 2010). Genetic polymorphism of the human *MUC5B* promoter sequence has been associated with diffuse panbronchiolitis and mucous hypersecretion (Kamio *et al.*, 2005). A single nucleotide polymorphism in the promoter region of the *MUC5B* gene has been linked to the development of familial interstitial pneumonia and sporadic idiopathic pulmonary fibrosis (Seibold *et al.*, 2011; Zhang *et al.*, 2011; Fingerlin *et al.*, 2013; Noth *et al.*, 2013; Stock *et al.*, 2013) and it has been suggested that this polymorphism might be associated with overexpression of *MUC5B* in the lung. More recently, a major function of MUC5B has emerged based on the findings of a unique *in vivo* study showing that MUC5B but not MUC5AC is essential for mucociliary clearance (Roy *et al.*, 2014).

We generated a mouse strain genetically deficient for Muc5b by deleting exons 12 and 13 of the 49 exons of the gene, exon 31 being the large central exon that codes for the Ser/Thr/Pro region (Desseyn, 2009). Here we report that no homozygous mice deficient for Muc5b were obtained while heterozygous mice were viable and fertile. Mice with Muc5b haplo-insufficiency displayed early lung inflammation that may lead to respiratory distress syndrome. In view of the embryolethality of full gene deletion, lung-restricted Muc5b-deficient mice (homozygous and heterozygous) were generated which show abnormalities of bronchial structure that may also lead to respiratory distress syndrome.

## RESULTS

### Absence of Muc5b is embryolethal

A targeting construct was developed to flank exons 12 and 13 of *Muc5b* gene by loxP sites (Fig. S1 and Fig. S2) located at the 5’ part of the gene, upstream of the large exon encoding the Ser/Thr/Pro region. Mice with the floxed *Muc5b* allele were intercrossed with the Cre deleter transgenic line MeuCre40. Mice carrying the Cre transgene and the Muc5b-floxed allele were backcrossed with C57BL/6 wild-type (WT) mice and their progeny with the Muc5b-floxed allele but without the Cre transgene were retained and studied. Muc5b^ko/+^ mice were fertile. Body mass was identical between Muc5b^ko/+^ and control WT mice (Muc5b^+/+^). Analysis of the progeny from 19 intercrosses of heterozygous Muc5b^ko/+^ mice (41 litters, 292 mice) was not consistent with Mendelian ratios as we observed 80 WT (27.4%) mice, 212 Muc5b^ko/+^ (72.6%) mice and no Muc5b^ko/ko^ progeny (*P*<0.0001). No homozygous embryos or resorption sites from E8 until birth (six pregnant mothers) were found suggesting that the complete tissue-disruption of both Muc5b alleles was embryolethal at a very early stage of embryogenesis. We then investigated the pulmonary phenotypes of the heterozygous systemic Muc5b^ko/+^ mice and lung specific Muc5b^ko/ko^.

### Some adult Muc5b^ko/+^ mice develop severe respiratory distress and abnormal lung histology

Of 63 Muc5b^ko/+^ mice analyzed, 12 (19%) displayed severe respiratory distress syndrome between 12 and 22 weeks of age including hunched posture, reduced locomotor activity, polypnoea (Video 1), a thumping respiration, squeaks and discreet cough (Video 2) while neither WT mice nor Muc5b-floxed mice displayed this pulmonary phenotype. Muc5b^ko/+^ mice developing respiratory distress displayed a dramatically abnormal lung morphology as illustrated in Fig. 1. Lung sections were stained with Mason’s trichrome stain, which readily identifies for Muc5b^ko/+^ mice deposition of collagen (blue color), a feature of pulmonary fibrosis. We also noted fibrin deposition, epithelial hyperplasia, interstitial inflammation and collapsed alveoli in comparison to WT mice (Fig. 1). Rarefaction of interstitial vessels was observed, as visualized by a reduction in CD31 endothelial positive cells (Fig. 1). The inflammatory cell infiltrate was also increased in Muc5b^ko/+^ mice, around the bronchi and vessels and was composed of both mononuclear and polynuclear cells (Fig. 1). Club cells (formerly Clara cells) were cuboidal shaped in Muc5b^ko/+^ mice showing bronchial metaplasia and hyperplasia (Fig. 2A). Increased Muc5b staining was observed in the bronchi and bronchioles of adult mice suffering from respiratory distress in comparison to WT control mice (Fig. 2B). Obstructive Muc5b positive material was occasionally observed in the lumen of the bronchioles in Muc5b^ko/+^ mice. In WT mice, Club cell secretory protein (CCSP) was present throughout the cytoplasm, whereas it was limited to the apical portion of Club cells in Muc5b^ko/+^ mice, consistent with a morphological change in the Club cells (Fig. 2C) in agreement with Boucherat *et al*. (Boucherat *et al.*, 2012). Total cell number of bronchial epithelium was increased (*P*=0.008; Fig. 2D) in Muc5b^ko/+^ mice with respiratory distress in agreement with the higher production of the mucin observed by immunohistochemistry. Bronchi of Muc5b^ko/+^ mice exhibited a decrease of CCSP-positive Club cells (*P*=0.03; Fig. 2D) with an increase of Muc5b-positive Club cells for Muc5b^ko/+^ mice with respiratory distress (*P*=0.03). No modification of the number of acetylated-tubulin (ACT) positive ciliated cells were observed (Fig. 2C and Fig. 2D). We then assessed the number of proliferative cells by immunofluorescence using anti-PCNA antibodies; this was significantly higher in Muc5b^ko/+^ mice than in Muc5b^+/+^ mice (*P*<0.0001; Fig. S3A and Fig. S3B) in agreement with Club cell hyperplasia.

**Figure 1.**
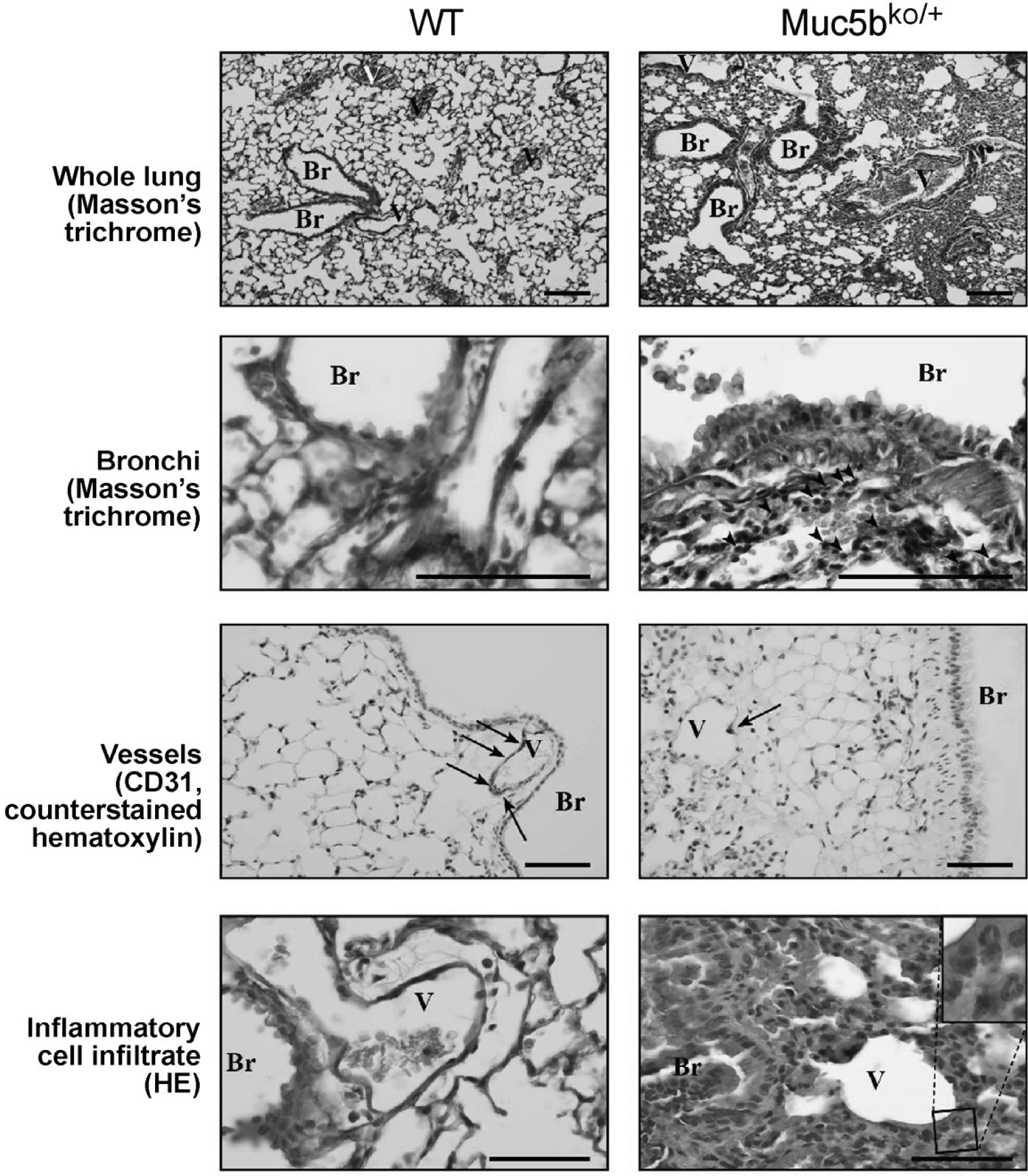
Representative histological analysis of adult lung tissue showing altered lung morphology of mice with respiratory distress. Histological sections at magnification x100 and x400. Inflammatory cells under the bronchial epithelium are indicated with arrowheads. CD31-positive cells are indicated with arrows. The inflammatory cell infiltrate is composed of both mononuclear and polynuclear cells, as shown at higher magnification in the insert. WT, wild-type. Br = bronchi; V = vessels.

**Figure 2.**
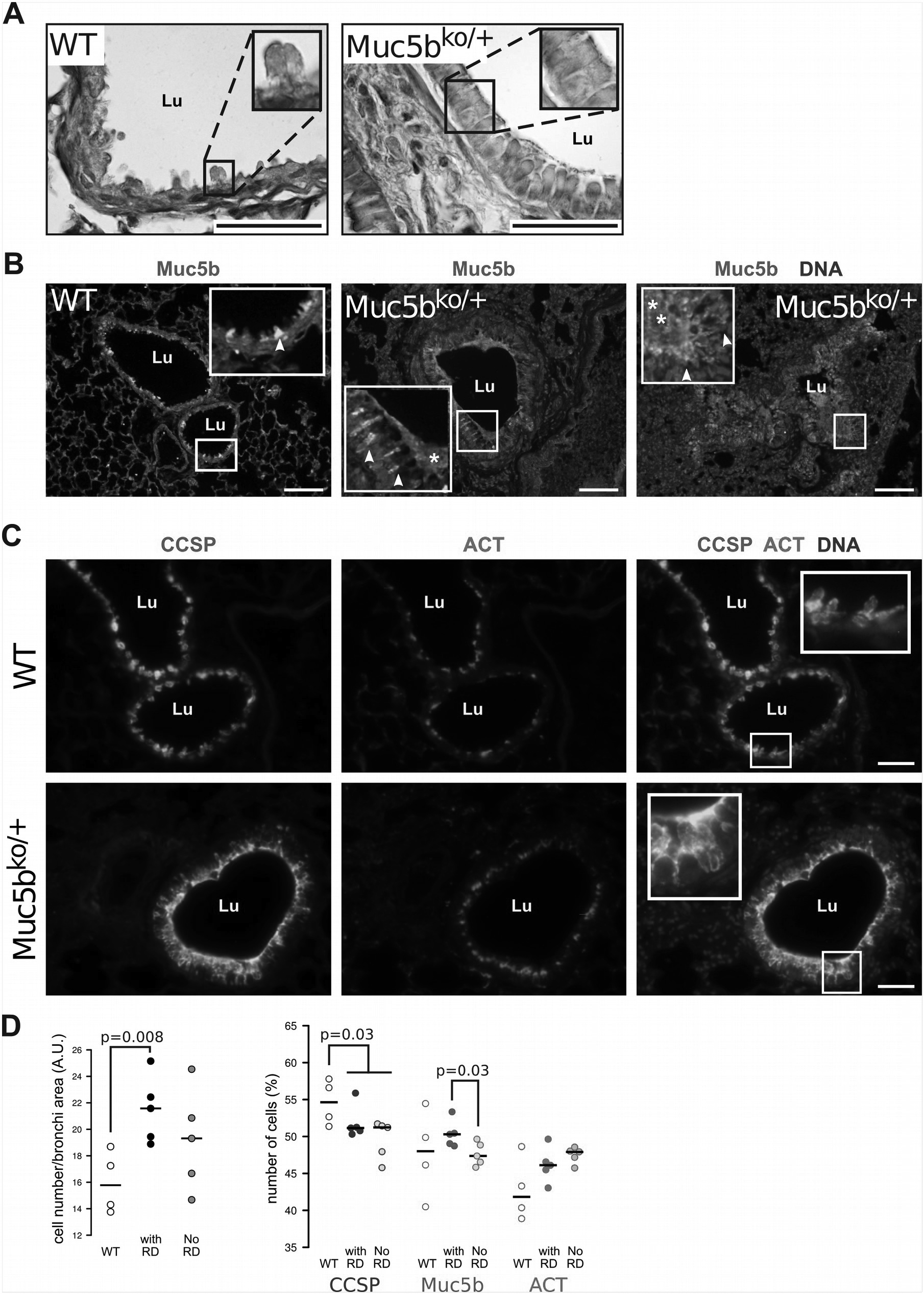
Histopathologic evidence of bronchial cell hyperplasia and goblet cell metaplasia. **(A)** Representative lung sections from adult wild-type (WT) and Muc5b^ko/+^ mice with respiratory distress stained with AB-PAS showing Club cells hyperplasia and metaplasia. Few goblet cells are visualized (AB-PAS+) in the WT mouse in comparison with the Muc5b^ko/+^ mouse. **(B)** Representative immunofluorescence of lung sections from wild-type (WT; n=1) with few goblet cells (Muc5b+) and Muc5b^ko/+^ mice (n=2) showing overproduction of Muc5b, goblet cell hyperplasia (arrow head) and mucus plug (*) in the lumen (Lu) of the bronchi in Muc5b^ko/+^. **(C)** Representative immunofluorescence of lung sections stained with anti-Club cell secretory protein (CCSP) and anti-acetylated-tubulin (ACT) in Muc5b^ko/+^ mice in comparison with WT mice. Muc5b^ko/+^ mice displayed goblet cell hyperplasia and Club cell metaplasia as revealed in the higher magnification in the inserts. Lu = lumen; Scale bar = 50 μm. **(D)** Bronchial cell density normalized to the bronchi area of four WT and five mice Muc5b^ko/+^ with respiratory distress (RD) and five Muc5b^ko/+^ mice without RD. **(E)** CCSP+ (blue), Muc5b+ (green) and ACT+ (red) epithelial cells of bronchi (%) from four WT, five Muc5b^ko/+^ with RD and five Muc5b^ko/+^ mice without RD. Six bronchi/mouse were assessed and analyzed using the Wilcoxon-Mann-Whitney test.

In WT mice, elastin fibers, which play a mechanical role in supporting and maintaining the lung tissue structure (Rocco *et al.*, 2009), ran longitudinally along the alveolar walls and were present at the tips of the alveolar septa of WT mice. By contrast, elastin fibers in the lungs of Muc5b^ko/+^ mice appeared disorganized and fragmented suggesting that the pulmonary tissue may be less elastic than in WT mice (Fig. S4A). Furthermore, occludin expression normally localized at the basolateral side of the airways epithelium was lost in epithelial cells of the bronchi or bronchioles of Muc5b^ko/+^ mice with an abnormal appearance of occludin expression in the alveolar epithelium (Fig. S4B).

To quantify histological changes including meta- and hyperplasia, inflammatory cell filtration of the parenchyma and fibrosis, lung sections were stained, coded, and then blindly scored (Madtes *et al.*, 1999). The mean score was significantly higher (*P*=0.008) in Muc5b^ko/+^ mice with respiratory distress than in wild-type mice (Fig. 3A). In Muc5b^ko/+^ mice, activated fibroblasts expressing α-smooth muscle actin (ASMA) secreting extracellular matrix component were increased around large (data not shown) and small airways (*P*=0.004; Fig. 3B and Fig. 3C) and also accumulated in the alveolar septae (Fig. 3D). Overall, these data suggest that deletion of one Muc5b allele may lead to severe respiratory distress, lung damage and affects lung histology with hyperplasia and metaplasia of bronchial cells and with fibrotic signs.

**Figure 3.**
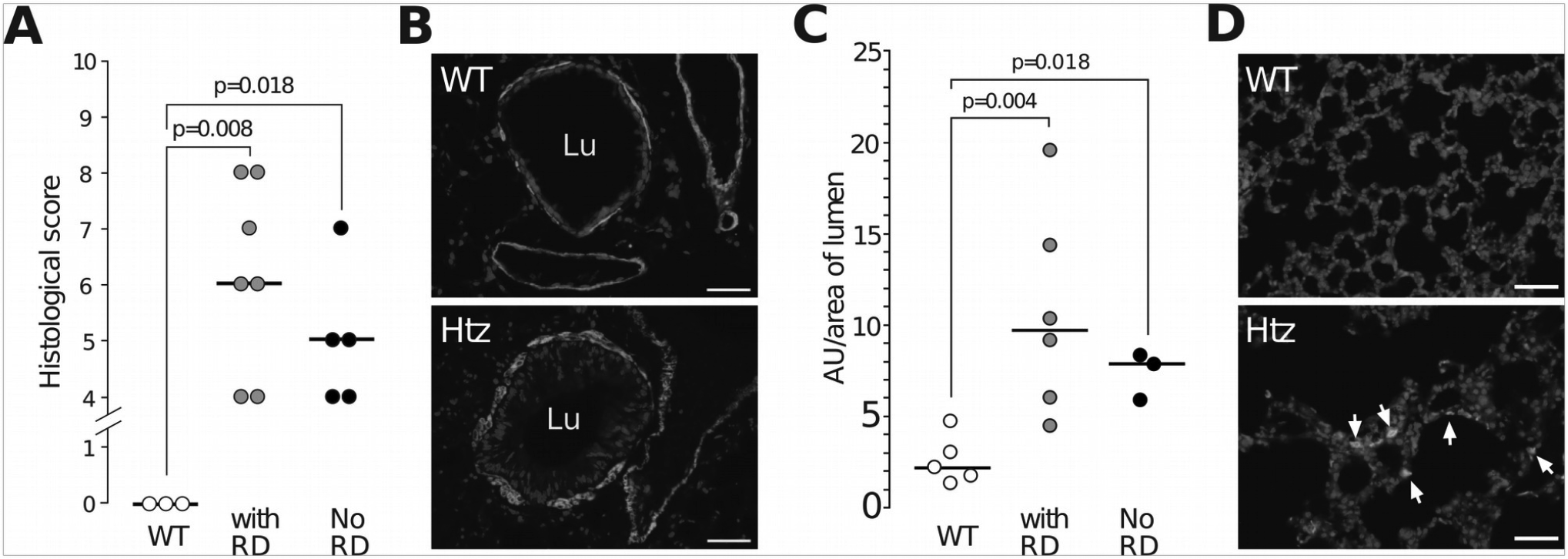
Increased histological score and expression of ASMA in the lungs of heterozygous adult mice. **(A)** Histological score of three wild-type (WT) mice, seven heterozygous mice with respiratory distress (RD) and five heterozygous mice without RD (no RD). **(B)** Representative immunofluorescence pictures of paraffin-embedded lung sections with anti-ASMA antibody. An increased in ASMA expression was observed around the bronchi of heterozygous (Htz; Muc5b^ko/+^) mice. **(C)** ASMA fluorescence was evaluated in the lungs of five WT (opened circles, 5 bronchi/mouse), six Htz mice with RD (4-5 bronchi/mouse) and three Htz mice without RD (4-5 bronchi/mouse). Data were analyzed using the Wilcoxon-Mann-Whitney test. **(D)** Representative immunofluorescence pictures with anti-ASMA antibody showing accumulation of ASMA in the alveolar space of Htz mice. Scale bar = 50μm.

### Adult Muc5b^ko/+^ mice without respiratory distress exhibit altered pulmonary function

To determine whether mice without signs of respiratory distress had abnormal lungs, we then studied 8/10- to 12/16-week-old mice. The lung morphology of 109 Muc5b^ko/+^ mice was analyzed macroscopically and 49 mice (45%) displayed an abnormal lung morphology characterized by the presence of abnormal grey areas and areas of necrosis and haemorrhage (Fig. S5A) but smaller than for mice with respiratory distress (Fig. S5B). Histological quantification showed that the lung score was significantly higher (*P*=0.018, Fig. 3A) for Muc5b^ko/+^ mice without respiratory distress than for WT mice. We then assessed the deposition of ASMA by immunohistochemistry followed by quantification of immunostaining. Increased deposition of ASMA was observed in the small bronchi of Muc5b^ko/+^ mice in comparison to WT mice (*P*=0.018; Fig. 3C) but the difference between mice with lung disease was not significant.

No macroscopic change was observed in other organs examined with the exception of the salivary glands since 13/51 (25%) Muc5b^ko/+^ mice exhibited atrophy of one salivary gland only (Fig. S5C). This organ was not investigated further.

To demonstrate functional abnormalities in the lungs from young Muc5b^ko/+^ mice, the lung mechanics of 10 WT/Muc5b^+/+^ and 10 Muc5b^ko/+^ 6-week-old male mice were analyzed using Flexivent at baseline and after metacholine administration. No significant baseline changes in dynamic resistance, Newtonian resistance, elastance, tissue damping and tissue elastance were observed in Muc5b^ko/+^ mice compared to WT mice (Fig. 4). We then administered metacholine, a smooth muscle agonist, to assess the effects of transient bronchoconstriction. An increase in doses of metacholine caused a decrease in maximum dynamic resistance in Muc5b^ko/+^ mice compared to WT mice, reflecting a decrease in level of constriction in the lungs (Fig. 4). Moreover, increasing the doses of metacholine caused a significant decrease in Newtonian resistance, which represents the resistance of the central airways in the constant phase model, elastance, tissue damping and tissue elastance in Muc5b^ko/+^ mice compared to WT mice.

**Figure 4.**
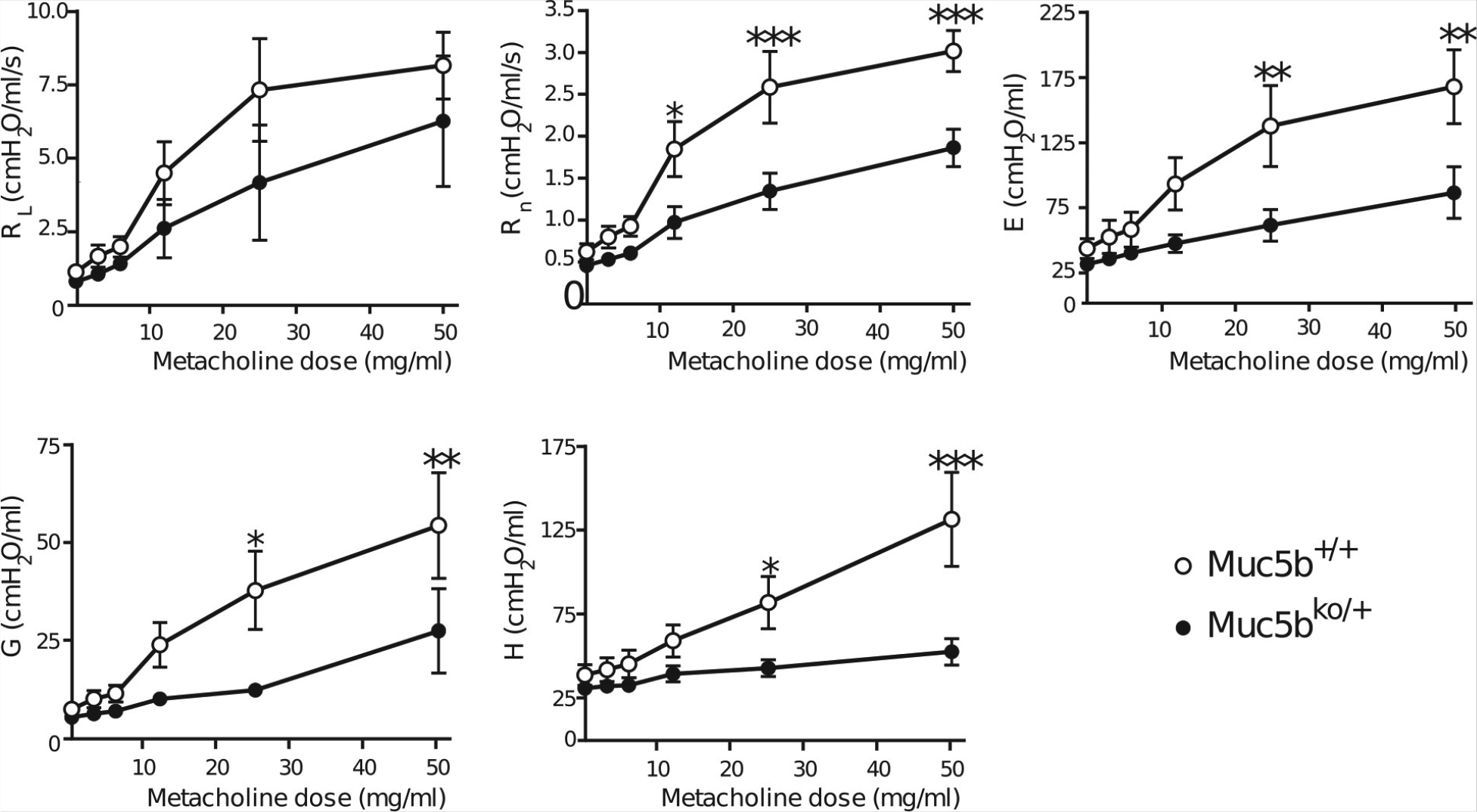
Muc5b^ko/+^ adult mice with no respiratory distress have altered lung function. The lung mechanics of eight wild-type (WT; opened circles) and eight Muc5b^ko/+^ (filled circles) mice were analyzed using Flexivent. No modification of baselines values of lung resistance (RL), Newtonian resistance (Rn), elastance (E), tissue damping (G) and tissue elastance (H) was observed in WT or Muc5b^ko/+^ mice. However, an increase in metacholine doses caused a decrease in RL, Rn, E, G and H in *Muc5b*^ko/+^ mice. * *P*=0.01; ** *P*=0.001; \*\*\**P*=0.0001. Data were analyzed using two-way analysis of variance.

### Adult Muc5b^ko/+^ mice show an increased lung inflammation in the absence of respiratory failure

As an inflammatory cell infiltrate was observed in both the lungs of 8-16-week-old Muc5b^ko/+^ mice (data not shown) and in older mice with lung disease, we assessed the levels of the pro-inflammatory chemoattractant chemokine CXCL1/KC in bronchoalveolar lavage (BAL) of adult WT and Muc5b^ko/+^ mice, which displayed no signs of respiratory insufficiency. While no CXCL1/KC was found in the 13 WT mice studied, CXCL1/KC was detectable in the BAL of 6/23 Muc5b^ko/+^ mice (26%; Fig. 5).

**Figure 5.**
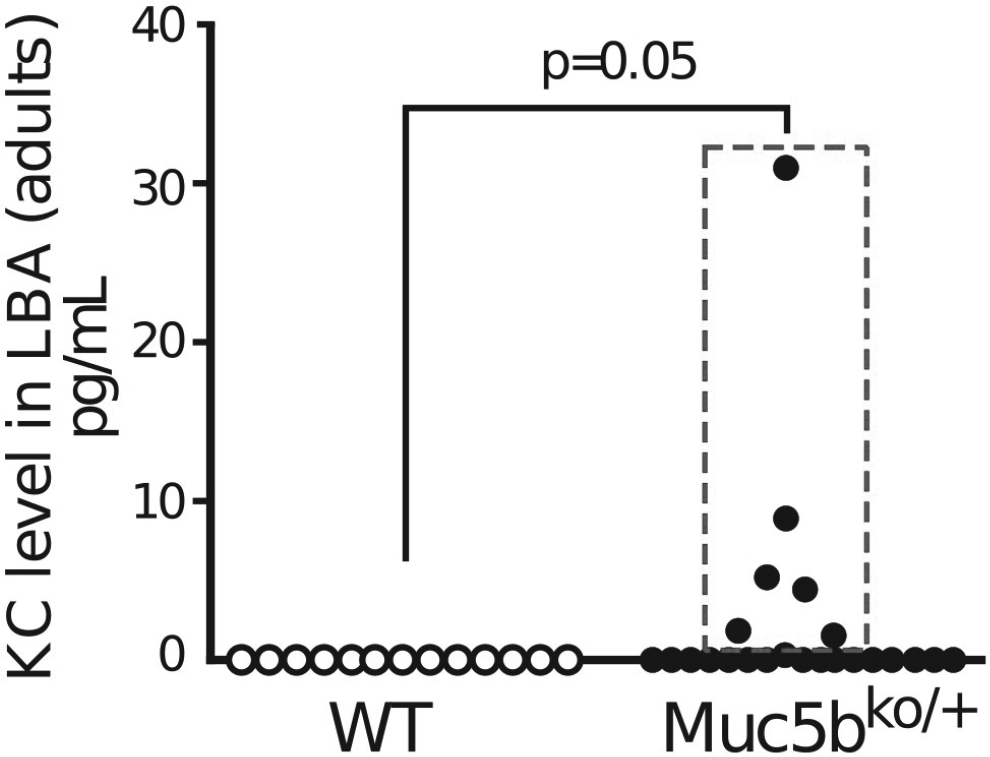
Inflammatory chemokine KC was elevated in lung of Muc5b^ko/+^ adult mice. Chemokine KC level was evaluated by ELISA in bronchoalveolar lavage (BAL) from 13 wild-type (WT) and 23 Muc5b^ko/+^ adult mice with no respiratory distress. The white circles emphasize 7/23 (30.4%) mice with detectable KC levels. Data were analyzed using Pearson’s Chi^2^ test.

### Pulmonary abnormalities appear early in life

To determine whether the lung defects in Muc5b^ko/+^ mice may appear early in life, we measured the CXCL1/KC level in whole lungs of 10 WT and 25 Muc5b^ko/+^ 2-day-old pups. A baseline CXCL1/KC level was detected in the lungs of WT mice which was significantly higher in Muc5b^ko/+^ mice as 6/25 (24%) Muc5b^ko/+^ mice displayed high levels of CXCL1/KC (*P*=0.03, Fig. 6A). Histology revealed that the lungs of 2-day-old Muc5b^ko/+^ pups had pathological changes at different levels (Fig. 6B). On whole lung sections, Muc5b^ko/+^ mice had a denser parenchyma, with condensed alveolae, congested vessels and an inflammatory infiltrate. The bronchial epithelium of Muc5b^ko/+^ mice was enlarged with an increased number of mucus-containing epithelial cells and numerous inflammatory cells in the lamina propria. Fibrin deposits, which are signs of tissue damage in lung injury, were found in the subepithelial region with an increased inflammatory cell infiltrate with abundant neutrophils in Muc5b^ko/+^ mice around the bronchi and vessels.

**Figure 6.**
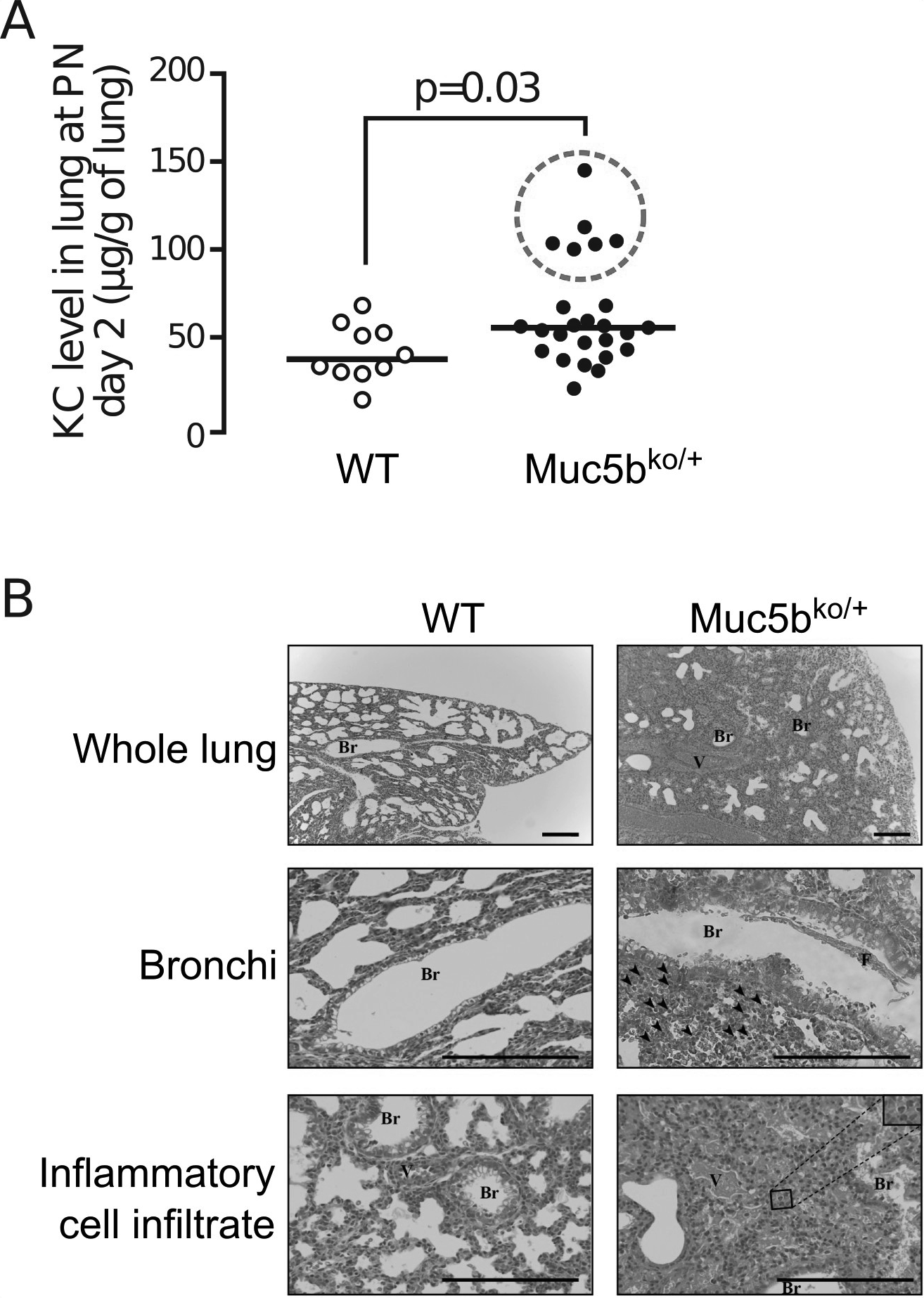
Inflammatory chemokine KC was elevated in lung of Muc5b^ko/+^ pup mice. **(A)** Chemokine KC level was evaluated by ELISA in the lungs of 10 wilt-type (WT) and 25 Muc5b^ko/+^ mice on post-natal day 2. The dotted red circle emphasizes 6/25 Muc5b^ko/+^ pups with high KC levels. Data were analyzed using the Wilcoxon-Mann-Whitney test. **(B)** Microscopic features of newborn mice on representative whole lung sections (magnification x100 and at higher magnification (x400)). Arrowheads denote inflammatory cells. Br and V indicate bronchi and vessels, respectively. F indicates fibrin deposits on the epithelial surface of the bronchial lumen. The higher magnification in the insert shows polymorphonuclear cells.

### Club cell-restricted Muc5b deficient mice have abnormal bronchi

Because Muc5b is not just expressed in the lung, the Muc5b mutation may cause an ectopic phenotype leading to a lethal phenotype. We then restricted the deletion of Muc5b in the lung using *CCSP* transgenic Cre mice crossed with Muc5b-floxed mice (Bertin *et al.*, 2005). CCSP, also referred to as CC10 and SCGB1A, is transcriptionally activated within the bronchi of neonatal mouse lungs starting at E16.5 (Reynolds *et al.*, 2002). Muc5b-floxed mice on one or two alleles were viable and fertile. No respiratory distress was observed in the 30 mice that were inspected by histology and which carried the CCSPCre transgene and no Muc5b-floxed allele (sacrificed between 40 and 50 weeks of age). Of the 124 mice studied and carrying the CCSPCre transgene and with at least one Muc5b-floxed allele, 27 (22%) were sacrificed as they showed signs of respiratory distress. Mice with conditional lung deletion of one or two Muc5b alleles, termed Muc5b^lung ko^, developed respiratory failure (Table 1; *P*<0.0001). The frequency of respiratory failure was increased when CCSPCre:Muc5b-floxed mice were between 20 and 30-weeks-old (Fig. S6) with a higher probability in mice that carried the two mutated alleles (65% *vs.* 35% for a single mutated allele.

**Table 1.**
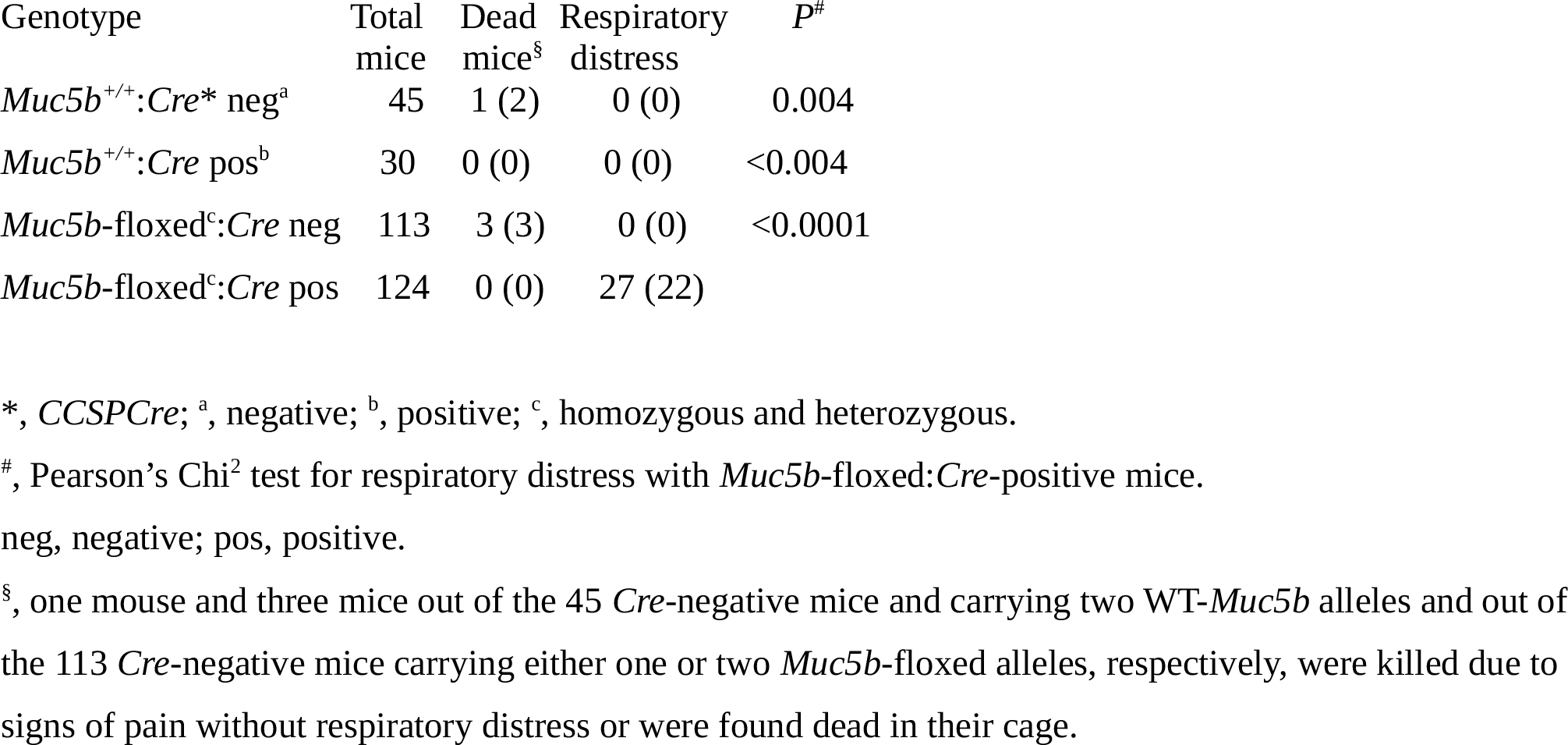
Respiratory distress before 50 weeks of age.

Lung histology revealed that the walls of the bronchi and bronchioles of 3-week-old Muc5b^lung ko^ mice but not adult mice was thinner than that of control floxed mice (CCSPCre-negative) with a flattened appearance of the ciliated cells (Fig. 7A). We then determined the total number of epithelial cells/mm^2^ of the cell wall of the bronchi of CCSP-positive cells and the number of ACT-positive cells by immunohistochemistry (Fig. S7). Reduced numbers of epithelial cells (*P*=0.007), CCSP-positive cells (*P*=0.02) and ACT-positive cells (*P*=0.02) were observed in Muc5b^lung ko^ mice (Fig. 7B and Fig. S7). As expected, low levels of Muc5b polypeptide were produced in Muc5b^lung ko^/CCSPCre mice when both alleles were floxed (heterozygous mice not shown) in contrast to homozygous floxed mice that were CCSPCre-negative (Fig. S7).

**Figure 7.**
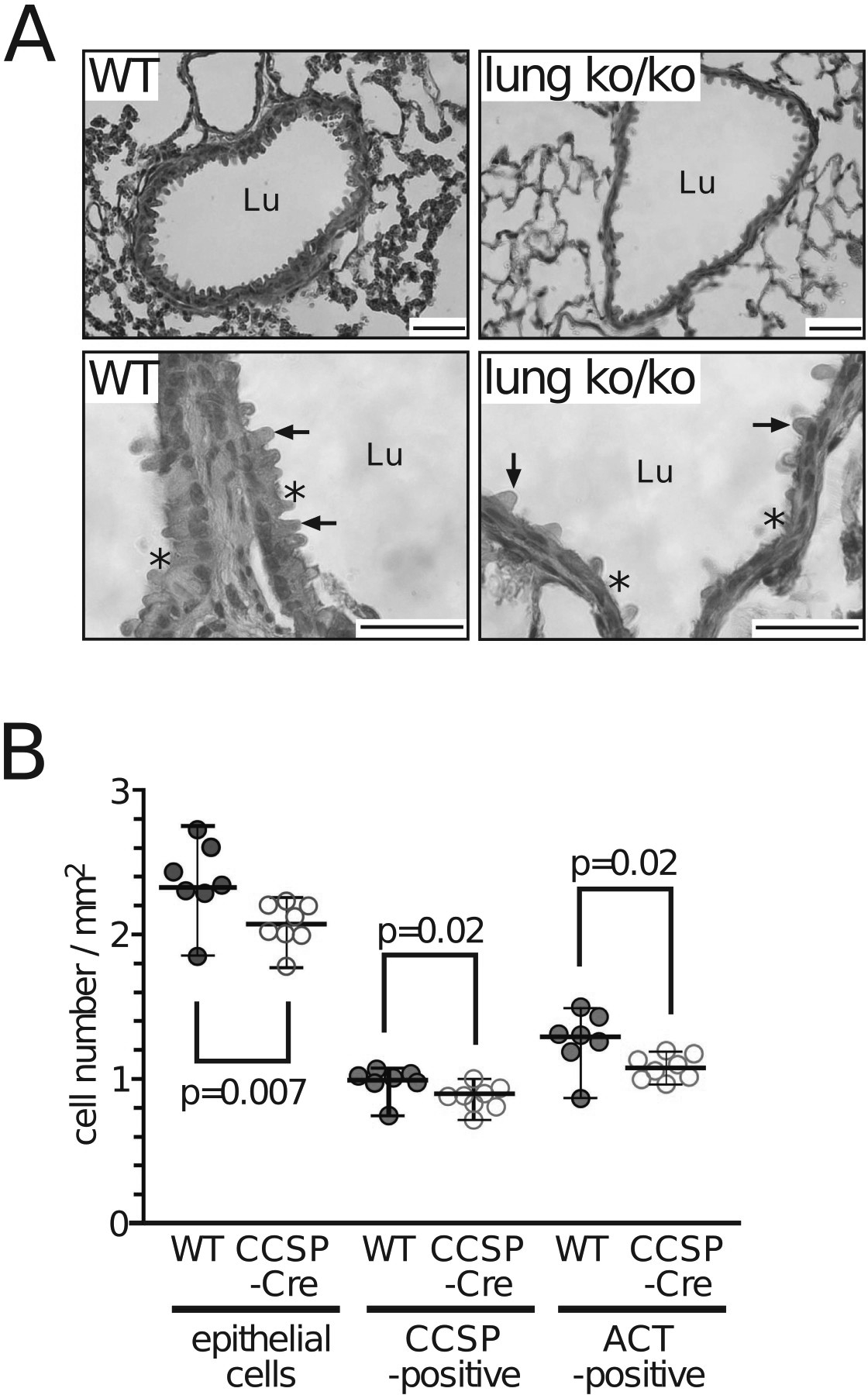
Histological analysis of Muc5b^lung ko/ko^ mice. **(A)** Representative histological analysis of paraffin-embedded lung sections of adult wild-type (WT) and Muc5b^lung ko/ko^ mice stained with hematoxylin-eosin at two magnifications. Muc5b^lung ko/ko^ mice displayed an abnormal morphology with depleted regions from epithelial cells. Stars indicate ciliated cells and arrows indicate Club cells. **(B)** Total cell number (blue), number of CCSP-positive (green) and ACT-positive (red) epithelial cell walls of bronchi from seven WT (filled circles) and eight Muc5b^lung ko^ (CCSP-Cre; opened circles) mice. Data were analyzed using the Wilcoxon-Mann-Whitney test.

## DISCUSSION

The two polymeric mucins MUC5AC and MUC5B represent the main gel-forming mucins in the mucus layer of the lung (Kesimer *et al.*, 2013). The precise function of each mucin is not well understood. Numerous reports link a dysregulation of the gel-forming mucin MUC5B in human lung diseases like asthma, cystic fibrosis, diffuse panbronchiolitis and familial and sporadic and interstitial pneumonia idiopathic pulmonary fibrosis (Kamio *et al.*, 2005; Rose & Voynow, 2006; Fahy & Dickey, 2010; Seibold *et al.*, 2011; Zhang *et al.*, 2011). In the current study, we characterized histological and functional parameters in lungs of mice that are genetically Muc5b-deficient. Alterations to lung, salivary glands and during embryogenesis are consistent with the distribution of Muc5b within wild-type mice (Roy *et al.*, 2014; Portal *et al.*, 2017*b*).

Bronchi are lined by a single columnar epithelium consisting of few goblet cells and ciliated and Club cells. Club cells can differentiate into ciliated and goblet cells. Club cell metaplasia/hyperplasia to goblet cell is a hallmark of lung damage in many airway diseases (see in Tompkins *et al*. (Tompkins *et al.*, 2009)). Goblet cell metaplasia and hyperplasia in bronchi of Muc5b^ko/+^ mice were evident in AB-PAS staining and immunohistochemical studies using anti-CCSP and anti-PCNA antibodies suggesting epithelial remodeling and hypertrophy in agreement with an increase production of Muc5b observed by immunohistochemistry (Fig. 2) and an increase of the inflammatory state. Invalidation of both alleles only in lung starting at E16.5, lead to minimal or no evidence of goblet cell metaplasia/hyperplasia. However, one fifth to one fourth lung-specific Muc5b-deficient mice exhibit also signs of respiratory distress in agreement with previous report showing the essential role of Muc5b for mucociliary clearance (Roy *et al.*, 2014; Livraghi-Butrico *et al.*, 2017). This suggests that the respiratory distress observed for conditional and all-tissues knockout mice comes from two distinct mechanisms and that Muc5b may play an important role in the Club cell differentiation.

Both Muc5b^ko/+^ with and without respiratory distress showed immune cell infiltrates, tissue injury and remodeling, differentiation of fibroblasts to myofibroblasts seen by ASMA expression and increased extracellular matrix deposition such as collagen which may explain the impaired ventilation and respiratory insufficiency we observed. These modifications are features of pulmonary fibrosis (Phan, 2002; Selman & Pardo, 2002; Scotton & Chambers, 2007; Li *et al.*, 2011; Seibold *et al.*, 2011; Wynn, 2011; Wuyts *et al.*, 2013; Camelo *et al.*, 2014) whereas airflow obstruction with progressive deterioration of lung function, mucus cell hyperplasia and immune cell infiltrates are hallmarks of chronic obstructive pulmonary disease (Jeffery, 1998). Whether or not Muc5b deficiency may lead to fibrosis was not directly demonstrated in our study and remains to be assessed.

Not all *Muc5b* mutated mice developed lung abnormalities, outlining a complex phenotype. Nineteen % of Muc5b^ko/+^ adult mice and 22% of Club cell-restricted Muc5b deficient adult mice that are 40 week-old or less suffered from severe respiratory distress syndrome. In the absence of respiratory distress, 45% of adult Muc5b^ko/+^ mice showed abnormal lung morphology and 26% showed abnormal levels of the pro-inflammatory chemokine CXCL1/KC in BAL consistent with inflammation in the lungs (Fig. 5). We have considered the possibility that abnormal pulmonary phenotype occurs early in life. At 2 days of age, 24% of Muc5b^ko/+^ pups already displayed abnormal high levels of the CXCL1/KC (Fig. 6A) although it is not possible to determine if the pups that displayed elevated CXCL1/KC levels correspond to mice which will develop respiratory distress at adulthood.

Agents responsible for the initiation of the lung phenotype are unknown. In lung fibrosis, it is believed that repeated lung injury could initiate inflammation cascades followed by overproduction of pro-fibrotic cytokines. Among the environmental triggers, viruses (Wuyts *et al.*, 2013) in our protected animal facility and which should not belong to the FELASA list (Mahler *et al.*, 2014) could be suspected as initiators of mouse fibrosis. Another possibility is mouse exposure to dust from bedding such as birch and hardwood. This has been reported in a case study to increase the risk of idiopathic pulmonary fibrosis in humans (Gustafson *et al.*, 2007). Further studies will be needed to investigate these hypotheses.

The development of a first Muc5b-deficient mouse line, term Muc5b^−/−^, has been reported (Roy *et al.*, 2014). The phenotype of Muc5b^−/−^ mice differs from that described in this current study. Muc5b^−/−^ mice showed impaired growth, survival (~40% at 12-months-old) and mucociliary clearance accompanied by abnormal breathing and material obstruction impeding the airflow in the upper airways. Inflammatory infiltrates and viable bacteria in the lung, especially streptococci and staphylococci, were also common in these mice. In this current study, we showed abnormal breathing and inflammatory infiltrates in Muc5b^ko/+^ mice and lung-deficient Muc5b mice but we never found any obstructing material during autopsy or culturable bacteria in the lung (data not shown) suggesting that the two different strategies used to mutate Muc5b and/or the animal environment may explain the variable phenotypes observed in the two models. Variable penetrance is common among mice deficient for the same gene and sometimes the phenotype may differ greatly as reported for example for genetically deficient mouse models for Nedd4-2, which has been shown to be essential for fetal and postnatal lung function (Boase *et al.*, 2011) and for CCSP-deficient mouse models (Zhang *et al.*, 1997; Reynolds *et al.*, 1999). The *Muc5b* gene has been mapped with the three other mucin genes *Muc2*, *Muc6* and *Muc5ac* to mouse chromosome 7 band F5 in a cluster of genes conserved in humans (Desseyn & Laine, 2003). The genomic organization and the deduced polypeptide sequence of the genes (amino- and carboxy-terminal regions), specially for *MUC5AC*, exhibit remarkable sequence similarities (Desseyn *et al.*, 1997, 1998, 2000; Buisine *et al.*, 1998). We cannot exclude cis-effects resulting from genetic modification itself of *Muc5b* and neighborhood effects to other adjacent unrelated genes or to *Muc5ac*, which is with *Muc5b* the other major gel-forming mucin in the lung.

Environmental conditions and variable genetic background in the two studies may explain in part the different predisposition of the two models to development of disease (DeMayo, 1999; Reynolds *et al.*, 1999). The two different gene-targeting strategies chosen to obtain Muc5b^−/−^ (team of CM Evans) and Muc5b^ko/+^ (and Muc5b^lung ko^, current investigation) mice may have yielded the different phenotypes. Because we deleted exons 12 and 13 and not the initiating transcription site, we cannot rule out that they are aberrant transcripts generating a hypomorph phenotype as previously reported for other genetically mouse deficient models (DeMayo, 1999) or truncated peptides translated from aberrant transcripts. We cannot rule out that other splicing events may have occurred too. Studies to understand the basis for differential phenotypes observed for the works on Muc5b-deficient mice should help to better understand the physiological functions of the mucin. The recent creation in our laboratory of a new conditional Muc5b-deficient mouse by floxing the last two exons of the gene (Portal *et al.*, 2017*a*) should also confirm in the near future that the lack of Muc5b leads to pulmonary distress without any bacterial infection of the lung.

## METHODS

### Generation of transgenic mice

The cloning of *Muc5b*, the construct of the targeting vector and transgenic mice generation are described in the online data supplement. Mice were maintained by breeding heterozygous mice after at least 6 backcrosses in C57BL/6 genetic background. In all experiments, mutated mice were compared with their WT littermates. The animal procedure followed in this study was in accordance with French Guidelines for the Care and Use of Laboratory Animals and with the guidelines of the European Union. The creation and use of the Muc5b floxed strain and progeny were approved by the French Biotechnologies Committee and registered under file number 5288.

### Histology and immunohistochemistry

Tissue fixation and coloration for histological and immunohistochemical analysis are described in the online data supplement. For immunohistochemistry, slides were incubated overnight at 4°C with antibodies against Muc5b (Valque *et al.*, 2011) (1:50), Club cell secretory protein (CCSP; 1:500; R42AP) (Ryerse *et al.*, 2001), occludin (1:100) (Invitrogen, 71-1500), proliferative cell nuclear antigen (PCNA; 1:100) (Abcam, PC10), α-smooth muscle actin (ASMA; 1:300) (Abcam, ab5694) or acetylated tubulin (ACT; 1:400) (Sigma, T7451) in PBS/1%BSA. For anti-PCNA and anti-ASMA antibodies, citrate buffer antigen retrieval was performed as described previously (Valque *et al.*, 2012). For anti-occludin antibodies, protease antigen retrieval was performed according to manufacturer’s instructions before BSA incubation. After three washes in PBS, slides were incubated with FITC-conjugated secondary antibodies (1:150) diluted in PBS/1%BSA for 2 h in a dark room at room temperature, rinsed three times in PBS and nuclei were counterstained with Hoescht 33258 (1:1000) for 5 min. For CD31 immunohistochemistry, chromogenic staining using horseradish peroxidase-3,3’ diaminobenzidine (HRP-DAB) staining was performed using an anti-CD31 antibody (Abcam, ab8364) diluted 1:50 according to the manufacturer’s instructions. For Muc5b, occludin and CD31 immunohistochemistry, lung sections of three control wild-type mice, seven 14-22-week-old Muc5b^ko/+^ mice with respiratory distress and five 10-16-week-old Muc5b^ko/+^ mice without respiratory distress were analyzed. Representative images of each labeling was illustrated in the figure.

Bronchi with similar diameters from five wild-type (WT) mice (5 bronchi/mouse), three Muc5b^ko/+^ mice (3-6 bronchi/mouse) without respiratory distress and six Muc5b^ko/+^ mice with respiratory distress (4-5 bronchi/mouse) were analyzed using an anti-ASMA antibody. ASMA immunofluorescence was measured using ImageJ software and expressed relative to the area of bronchial lumen.

Five to 7 bronchi/adult mouse with similar diameters from four WT, five Muc5b^ko/+^ with respiratory distress and five Muc5b^ko/+^ without respiratory distress were analyzed using anti-Muc5b, anti-CCSP and anti-ACT antibodies. Concerning lung-specific Muc5b knockout mice, bronchi with similar diameters from seven WT mice (5-7 bronchi/mouse) and from eight 3-week-old Muc5b floxed mice carrying the CCSPCre transgene (5-8 bronchi/mouse) were analyzed using anti-ACT and anti-CCSP antibodies. The area of the bronchial wall was determined using ImageJ software by subtracting the area of the bronchial lumen from the total area of the bronchus. Total epithelial cells from bronchi, Muc5b-immunopositive (goblet cells), CCSP-immunopositive cells (Club cells), ACT-immunopositive cells (ciliated cells) and unknown cells (neither ACT and CCSP-positive) were counted and expressed relative to the area of the epithelial cell wall of the bronchi.

The number of PCNA-immunopositive cells was counted by observing 10 randomly selected bronchi or bronchioles in lung sections from three WT and three heterozygous mice.

### Histological score

Lung sections from three control wild-type mice, seven 14-22-week-old h Muc5b^ko/+^ mice with respiratory distress and five 10-16-week-old Muc5b^ko/+^ mice without respiratory distress were scored blindly for inflammation and fibrosis from 0 to 11 (adapted from Madtes *et al.* (Madtes *et al.*, 1999)) and metaplasia and hyperplasia of the bronchi. Lung inflammation, metaplasia and hyperplasia of the bronchi was scored on H&E-stained lung sections on a scale of 0 to 3 for inflammation (0 = no inflammatory involvement; 1 = inflammatory cell infiltration of 3% to 29% of the parenchyma; 2 = inflammatory cell infiltration of 30% to 59% of the parenchyma; and 3 = inflammatory cell infiltration of 60% to 100% of the parenchyma), on a scale of 0 to 2 for metaplasia (0 : no metaplasia ; 1 : slight metaplasia of epithelial cells of the bronchus ; 2 : strong metaplasia) and on a scale of 0 to 2 for hyperplasia (0 : no hyperplasia ; 1 : slight hyperplasia of epithelial cells of the bronchus ; 2 : strong hyperplasia). Lung fibrosis was scored on trichrome-stained lung sections on a scale of 0 to 4 (0 = no increase in connective tissue; 1 = fine connective-tissue fibrils in less than 50% of the area occupied by inflammatory cells, without coarse collagen; 2 = fine fibrils in 50% to 100% of the same area, without coarse collagen; 3 = fine fibrils in 100% of the area, with coarse collagen bundles in 10% to 49% of the area; 4 = fine fibrils in 100% of the area, with coarse collagen in 50% to 100% of the area). Scores were established on 3 lobes (left lobe and right cranial and middle lobes) of lung. Three different sections of the lung (upper, middle and lower parts) were analyzed.

### Lung mechanics

Lung mechanics were assessed in Muc5b^ko/+^ mice and Muc5b^+/+^ brother-sister mice using Flexivent (Scireq, Montreal, Canada) as described in the supplemental data. The maximum dynamic resistance (R_L_), Newtonian resistance (Rn), elastance (E), tissue damping (G) and tissue elastance (E) were recorded before and after increasing doses of aerosolized metacholine.

### Measurement of chemokine KC in bronchoalveolar lavage

Bronchoalveolar lavage (BAL) was performed prior to sacrifice by two consecutive injections (500 μL and 1 mL) of PBS through a tracheal canula. Lavage fluid was centrifuged at 1000 *x g* for 10 min and the supernatant was stored at −80°C until use. Chemokine KC was measured using an ELISA kit (R&D systems, DY453) according to the manufacturer’s instructions. Absorbance at 450 nm was determined using a microplate reader. Fifty μL of the BAL supernatant was used to evaluate the KC level of adult mice. To measure the KC level in lung at post-natal day 2, newborn mice were sacrificed, lungs were removed and homogenized in PBS. The homogenate was then centrifuged at 1000 *x g* for 10 min and 50 μL of the supernatant was used to quantify the KC level. Lungs used for BAL were not used for other investigations.

### Statistical analysis

For histological score, ASMA quantification, KC level quantification in PCNA-positive cells, cell number quantification (ACT-, CCSP-positive cells and total epithelial cells of bronchi), the graphs show the median value. The Wilcoxon-Mann-Whitney and Pearson’s Chi^2^ tests were performed using StatXact 6.0 (Cytel Studio, Cambridge, MA) to compare unpaired data. For the lung mechanics, all results are expressed as mean ± standard error of the mean. Two-way analysis of variance was performed using GraphPad Prism software (La Jolla, USA) and was used to analyze lung parameters. A *P* value of ≤0.05 was considered statistically significant.

## COMPLIANCE AND ETHICS

The authors declare that they have no conflict of interest. All experiments conformed with the Helsinki Declaration of 1975 (as revised in 2008) concerning Animal Rights.

## ACKNOWLEDGEMENTS

We thank M. Holzenberger (Inserm UMRS 938, Paris, France) for the MeuCre40 mouse strain, M. Tauc (CNRS FRE 3093, Nice, France) for the CCSPCre mouse strain, C. Goujet-Zalc (CNRS, SEAT UPS44, Villejuif, France) for the generation of Tg mice, J.S. Ryerse (Dept. of Pathology, St Louis University, MO, USA) for the anti-CCSP antibody, M.H. Gevaert and R.M. Siminski (Service Commun-Morphologie Cellulaire, Univ. Lille, France) for slides, J. Devassine and D. Taillieu from the EOPS animal facility (Univ. Lille, France) for mouse colony management and P. Roussel for critical reading of the manuscript and useful discussions. This study was supported in part by the French Cystic Fibrosis Association – Vaincre la Mucoviscidose.

## Methods

### Generation of transgenic mice

#### Muc5b *gene cloning*

The mouse *Muc5b* gene has been mapped on chromosome 7 band F5 in a cluster of genes conserved between humans and mice (Desseyn & Laine, 2003). The two exonic oligonuccleotides 5’-GACGTCTTCCGCTTCCCTGGCCT-3’ and 5’-TCTTCATTCCACAGGAAGGT-3’, respectively, were designed from the *Muc5b* gene (Chen et al., 2001). These two primers flank intron 3 and were used to amplify genomic DNA extracted from mouse embryonic stem (ES) cells by PCR. A genomic sequence of 437 bp was cloned and sequenced showing that it contained an intron of 144 bp belonging to *Muc5b*. The 437 bp insert was used as a probe to screen a mouse 129Sv bacterial artificial chromosome (BAC) clone library (Incyte Genomics). Two positive BAC clones were identified. One BAC clone was purchased and was shown to contain the full genomic sequence of *Muc5b* (data not shown). Both human and mouse *Muc5b* genes consist of 49 exons, with exon 31 being the largest (10.7 kb in humans (Desseyn *et al.*, 1997) and encoding the *O*-glycosylated regions of mucin.

#### Targeting vector design

The general 3-loxP strategy used to invalidate *Muc5b* is summarized in Figure E1A and B. Deletion of exons 12 and 13 should lead to a frameshift introducing a premature stop codon. The loxP sequence (34 bp) was subcloned into the pKS+ plasmid (Stratagene) between the unique *Eco*RI and *Xho*I restriction sites. The *Muc5b* targeting construct utilized 5.5 kb and 3.9 kb genomic fragments of the 5’-end of *Muc5b* that flanked the *Nhe*I restriction site in intron 13 as the left and right arms, respectively. A unique loxP site was introduced in intron 11 and a blunted-*Xho*I*-Xho*I loxP-flanked neomycin expression phosphotransferase (NEO) cassette (Howe *et al.*, 2006) was inserted in the blunted-*Nhe*I site of intron 13. A recombinant plasmid carrying three loxP sites with the same orientation was selected. Deletion of the genomic region carrying exons 12 and 13 with Cre recombinase should introduce a frameshift leading to a premature stop codon. The correct orientation of the three loxP sites was verified by transforming bacteria carrying the plasmid construct with a plasmid encoding Cre recombinase (New England Biolabs; Figure E2). Plasmid DNA was then linearized using the unique *Eco*RV restriction site located within intron 17 and analyzed on a 0.6% agarose gel.

#### Transgenic mice

The targeting vector was linearized with *Eco*RV and digested using the *Pvu*I restriction enzyme found twice in the pKS plasmid in order to excise the plasmid insert. The 10.6 kb insert was electroporated in CK35 ES cells (SEAT, Villejuif, France). The genomic DNA of ES cells and mouse tissues was extracted, purified and 10 μg of DNA was digested with *Kpn*I and electrophorezed on a 0.8% agarose gel. DNA was transferred onto a nylon membrane (Roche Applied Science) and probed with the labeled PCR product described above. Probe hybridization was performed at 42°C overnight with shaking. Detection was carried out by chemiluminescence using an anti-digoxigenin Fab antibody and CDP-star according the manufacturer’s instructions (Roche Applied Science). The 689 bp *Muc5b* probe was obtained by PCR amplification using the two oligonuccleotides 5’-TGGGCATCCCACTTGCTG-3’ (forward) and 5’-GTAGAGAGGGTCAACTGATGC-3’ (reverse) and labelled with digoxigenin (DIG)-labelled 11-dUTP (Roche Applied Science). One positive ES cell clone was obtained (Figure E1C) and microinjected.

Chimeric male mice were obtained and intercrossed with C57BL/6 wild-type (WT) mice purchased from Charles River, France. The resulting offspring with the mutated *Muc5b* locus carrying three loxP sites and their progeny were kept in a specific-pathogen free animal facility. DNA extracted from tail biopsies was analyzed by PCR amplification using the two oligonucleotides 5‘-GAGAGGCCTCCACTCTTTCTCCAAGC-3’ (P1; forward) and 5’-CCAAATGTGCATGGCGTGTAAATGAC-3’ (P2; reverse) that flanked the loxP site located within intron 11. Heterozygous mice were intercrossed with the MeuCre40 strain (C57BL/6 genetic background).

MeuCre40 genotyping was performed as described elsewhere (Leneuve *et al.*, 2003). Pups carrying both the mutated *Muc5b* locus and the Cre sequence were bred with WT C57BL/6 mice. After two generations, mice without the Cre sequence but carrying the *Muc5b* allele deleted for exons 12 and 13 (Muc5b^ko/+^) without the NEO cassette were kept and intercrossed. The Muc5b genotype of mice was determined by PCR amplification using the oligonucleotides P1 coupled to the oligonucleotide 5’-GAGAAGAAAGTCCCCGCCCAGTGTTT-3’ (P3; reverse) to amplify the knock-out (KO) allele. Deletion of the selective cassette was performed using the primer P2 coupled to the specific NEO cassette oligonucleotide 5’-TGTTGTGCCCAGTCATAGCCGAATAG-3’ (P4; forward). Genotyping was confirmed by Southern blotting with the external 5’ probe and an internal probe using the two oligonucleotides 5’-GTGGAGAGGCTATTCGGCTATG-3’ and 5’-CTCTTCAGCAATATCACGGGTAG-3’ amplifying a 648 bp NEO nucleotide sequence. Muc5b^ko/+^ heterozygous mice were backcrossed for at least five generations into the C57BL/6 genetic background. Muc5b floxed mice that do not carry the MeuCre40 transgene were also bred with CCSPCre mice to obtain lung-specific Muc5b-deficient mice (Bertin *et al.*, 2005). The genotype of mice carrying CCSPCre was determined by PCR using the two oligonucleotides used to genotype the MeuCre40 transgene.

### Histology and immunohistochemistry

Mice were anesthetized by injection of 200 μL pentobarbital. The lungs and salivary glands were gently removed, rinsed in PBS and fixed in 4% paraformaldehyde in PBS for 20 h. Formalin-fixed tissues were dehydrated through a series of increasing ethanol washes and embedded in paraffin. Paraffin blocks were brought to room temperature and sectioned on a rotary microtome. Five μm thick sections were floated onto water at 40°C before being transferred to Superfrost/Plus microscope slides (Thermo Scientific). Sections of paraffin-embedded lung tissue were dewaxed with xylene, rehydrated through a series of decreasing ethanol washes and stained with haematoxylin-eosin (HE), alcian blue-periodic acid Schiff (AB-PAS) and Masson’s trichrome for microscopic examination. For the elastin fiber study, sections were stained with orcein (Sigma-Aldrich). Tissue sections were analyzed blindly by two different pathologists unaware of the genotypes on a motorized Z-axis microscope (BX 61 Olympus, Tokyo, Japan), using epi-fluorescent light. Microscope pictures were obtained with a digital camera ColorView III using Olympus-SIS Cell F software (Olympus, Tokyo, Japan).

For immunohistochemistry, sections of paraffin-embedded lung tissue were dewaxed with xylene and rehydrated through a series of decreasing ethanol washes and rinsed three times in PBS. To block non-specific binding, slides were incubated with 1% bovine serum albumin (BSA) in PBS for 45 min. The immunolabeled sections were dried and mounted with Mowiol mounting medium and stored at 4°C. Images were acquired and were minimally processed as described previously (Gouyer *et al.*, 2010).

### Lung mechanics

Lung mechanics were assessed in Muc5b^ko/+^ mice and Muc5b^+/+^ brother-sister mice using Flexivent (Scireq, Montreal, Canada) as follows. Six-week-old mice were anesthetized by intraperitoneal injection of medetomidine (5 mL/kg; Pfizer, Paris, France) and 10% ketamine (Merial, Lyon, France), paralyzed by intraperitoneal injection of 1% pancuronium bromide (5 mL/kg; Organon) and immediately intubated with an 18-gauge catheter, followed by mechanical ventilation. Respiratory frequency was set at 150 breaths/min with a tidal volume of 0.2 mL and a positive-end expiratory pressure of 2 mL H_2_O was applied. Mice were exposed to nebulised PBS followed by increasing concentrations of nebulised metacholine (3-50 mg/mL in PBS; Sigma-Aldrich) using an ultrasonic nebuliser (Aeroneb, Aerogen, Galway, Ireland). For each dose, 10 cycles of nebulisation and measurements were performed. Nebulisation was conducted during the first cycle and consisted of 20 puffs per 10 s, with each puff of aerosol delivery lasting 10 ms. For each cycle, measurements were obtained for 15 s followed by ventilation for 5 s. The maximum dynamic resistance (R_L_), Newtonian resistance (Rn), elastance (E), tissue damping (G) and tissue elastance (E) were recorded before and after increasing doses of aerosolised metacholine.

---

**Figure S1.**
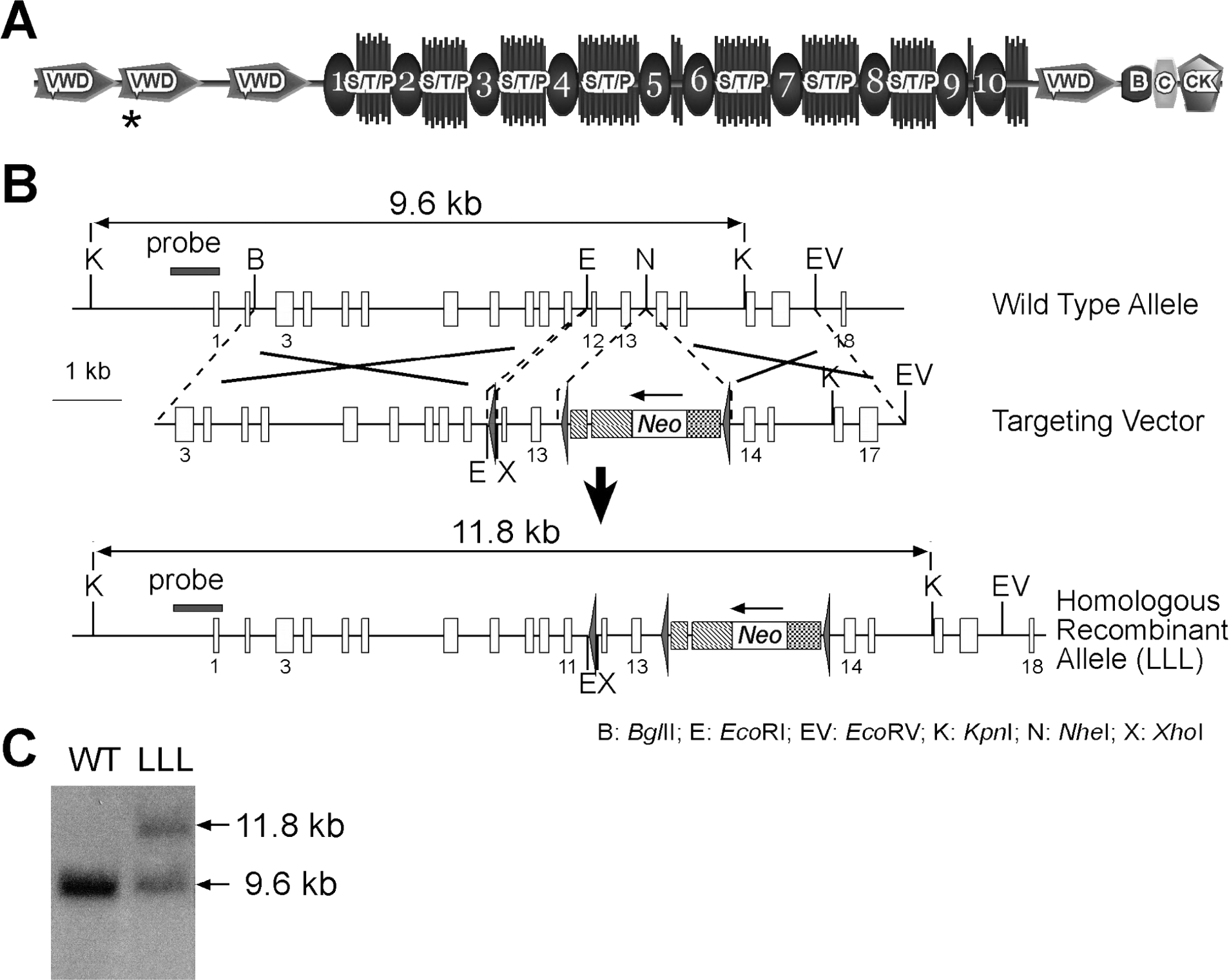
Targeted disruption of the *Muc5b* gene. **(A)** Schematic domain structure of mouse Muc5b protein. Mouse Muc5b protein is composed of amino- and carboxy-terminal regions that share similar domains with von-Willebrand factor (vW-B, -C, -CK and -D). The central part of Muc5b contains large regions enriched in Ser, Thr and Pro (S/T/P), which are extensively substituted with *O*-glycans. A highly conserved domain (oval blue) named CYS domain is found 10 times in Muc5b and is linked to or interrupts the Ser/Thr/Pro regions. The star indicates the targeted region used to invalidate *Muc5b*. **(B)** Structure and partial restriction enzyme map of the genomic segment of *Muc5b* used to generate the targeting vector. Empty boxes indicate exons and some of them are numbered. In the targeting vector, exons 12 and 13 are flanked by loxP sites and a NEO cassette flanked by a third loxP site was inserted into intron 13, in the opposite transcriptional orientation to that of *Muc5b*, as indicated by the arrow. **(C)** Southern blot analysis of DNA isolated from a control ES clone (WT) and the positive ES cell clone (LLL). DNA was digested with *Kpn*I and hybridised with the 5’ probe (indicated by a grey box in (B)). The 9.6 and 11.8 kb bands correspond to the WT and mutant allele (LLL allele), respectively.

**Figure S2.**
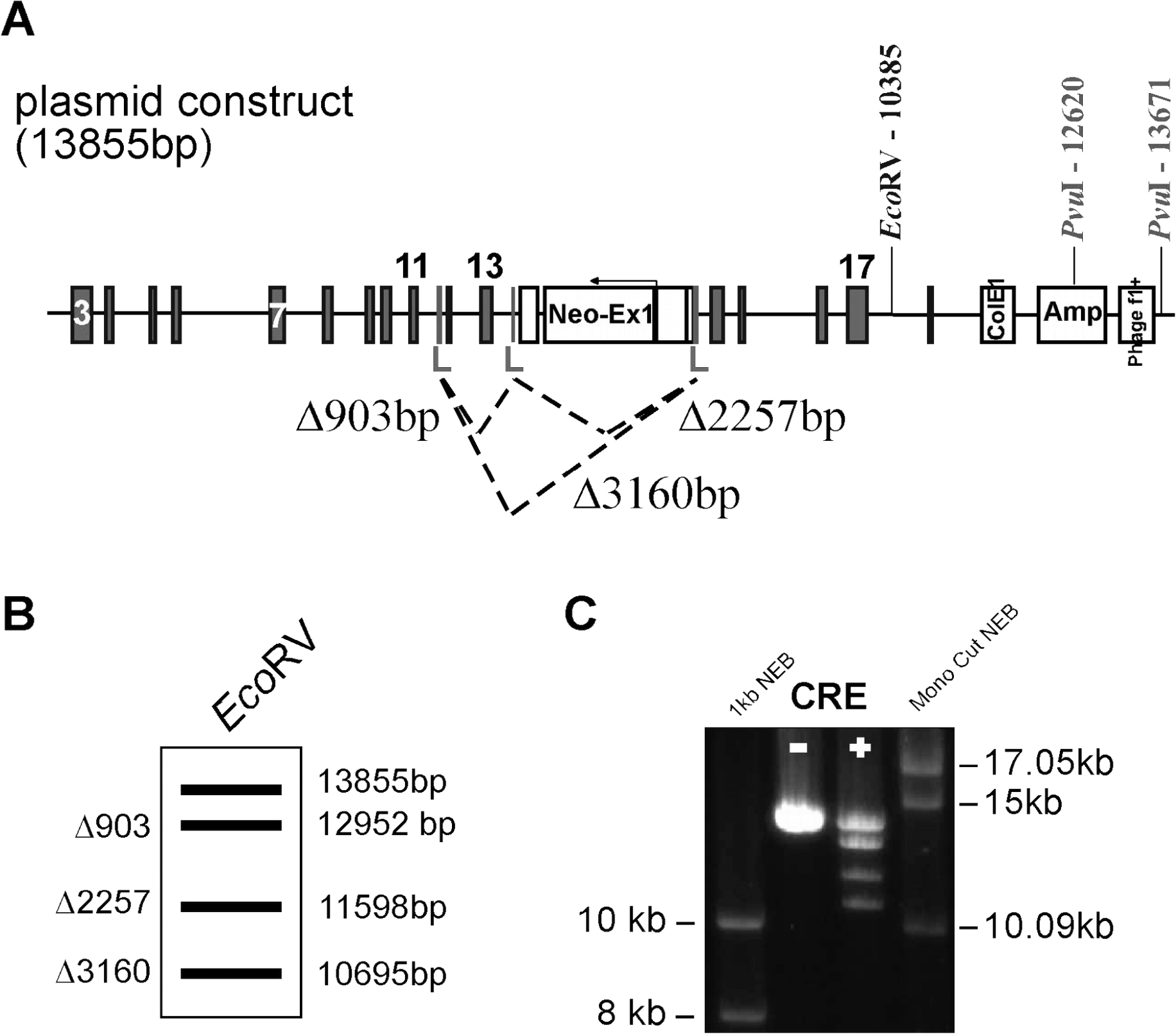
The three loxP sites are in the same orientation. **(A)** Schematic representation of the targeting vector. The three deletion 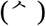 events are represented. **(B)** Predicted profile of *Eco*RV-linearised plasmid on agarose gel after Cre recombination if the three loxP sites have the same orientation. **(C)** Following incubation (+) of the circular plasmid construct with Cre recombinase (New England Biolabs) for 3 h, the DNA was digested with *Eco*RV and analyzed on a 0.6% agarose gel in comparison with the plasmid construct without Cre recombinase (−). Plasmid DNA with Cre (+) showed one band corresponding to the unrecombined DNA (13.9 kb) and three lower bands that correspond to the three possible recombination events depicted in **(A)**. The two DNA ladders are the 1 kb-ladder and the monocut-ladder from New England Biolabs.

**Figure S3.**
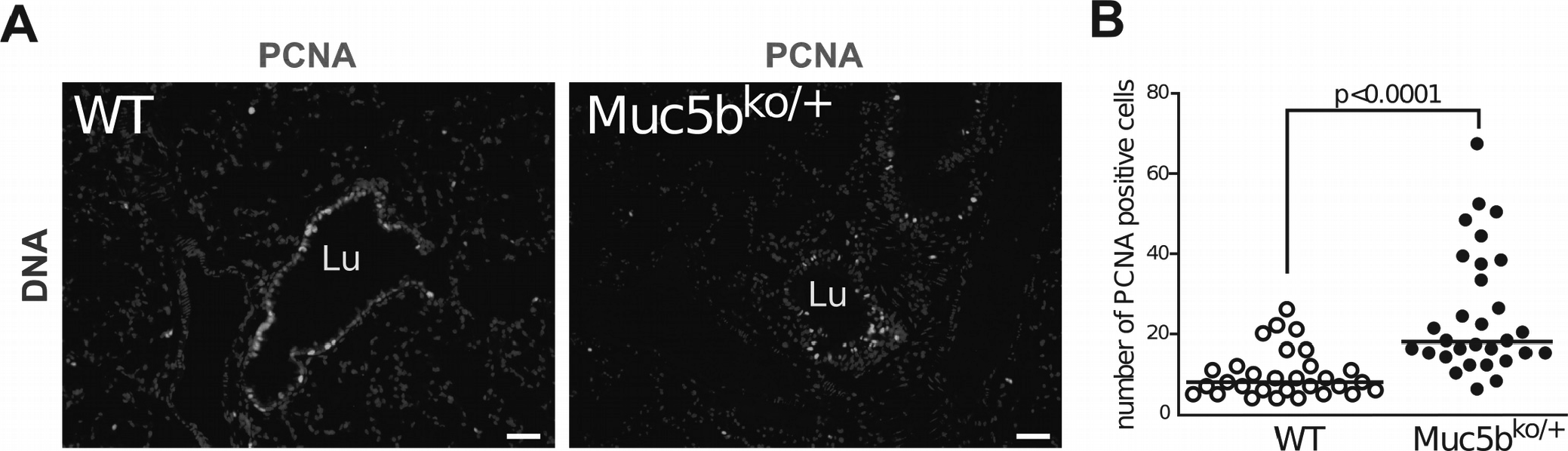
Increase cell proliferation in Muc5b^ko/+^ mice with respiratory distress. **(A)** Representative immunofluorescence pictures of paraffin-embedded lung section of a wild-type (WT) and a Muc5b^ko/+^ mouse stained with anti-proliferative cell nuclear antigen (PCNA) antibody. Lu = lumen; Scale bar = 50 μm. **(B)** The number of PCNA-positive cells was evaluated in the lungs of three WT (opened circles; 10 bronchi/mouse) and three Muc5b^ko/+^ (filled circles; 10 bronchi/mouse) mice. Data were analyzed using the Wilcoxon-Mann-Whitney test.

**Figure S4.**
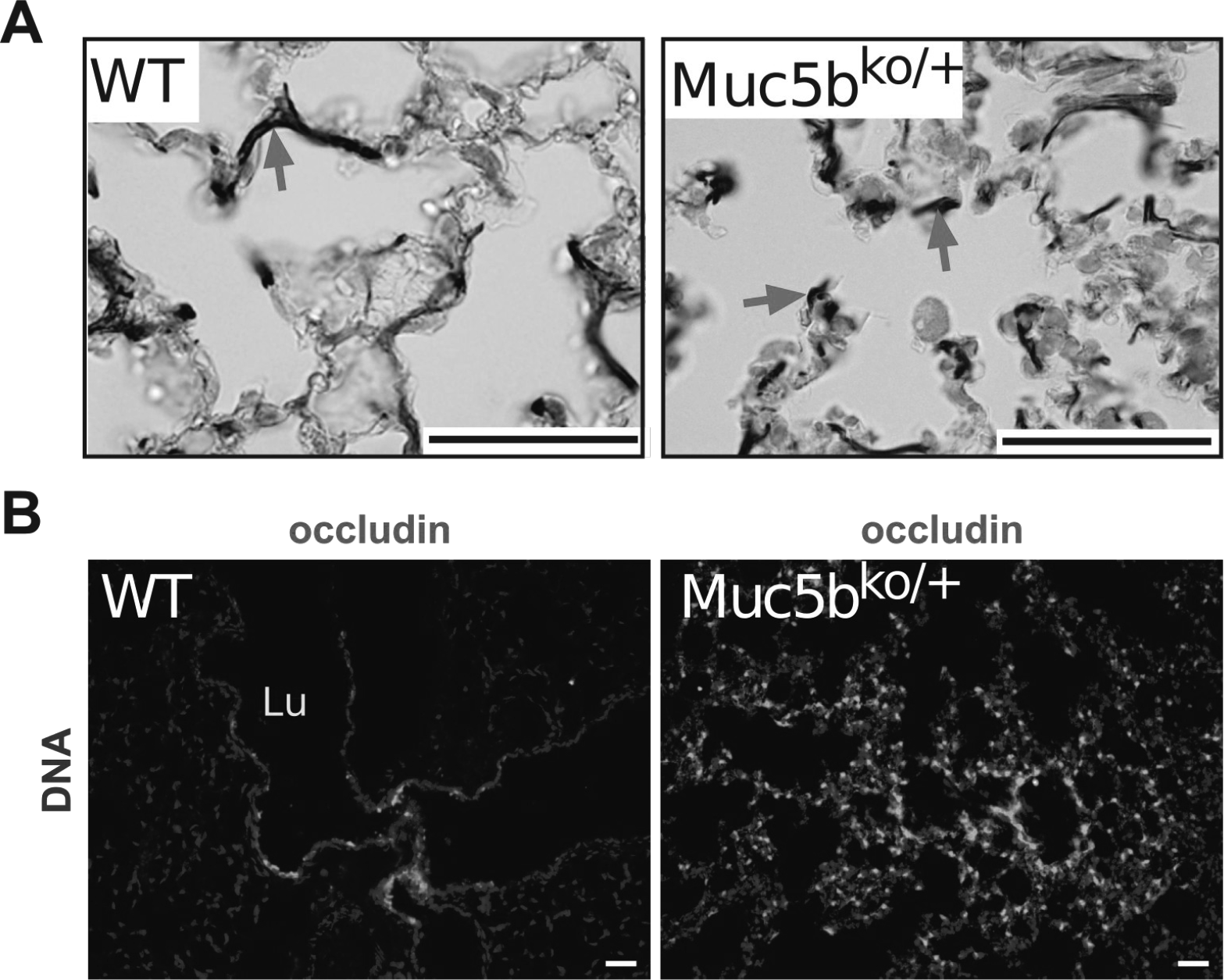
Disorganization of tight junctions of Muc5b^ko/+^ mice with respiratory distress. **(A)** Orcein staining showing disorganization of elastin fibers (red arrows) in Muc5b^ko/+^ mice. **(B)** Representative immunofluorescence pictures of paraffin-embedded lung sections stained with anti-occludin antibody. Lu = lumen; Scale bar = 50 μm.

**Figure S5.**
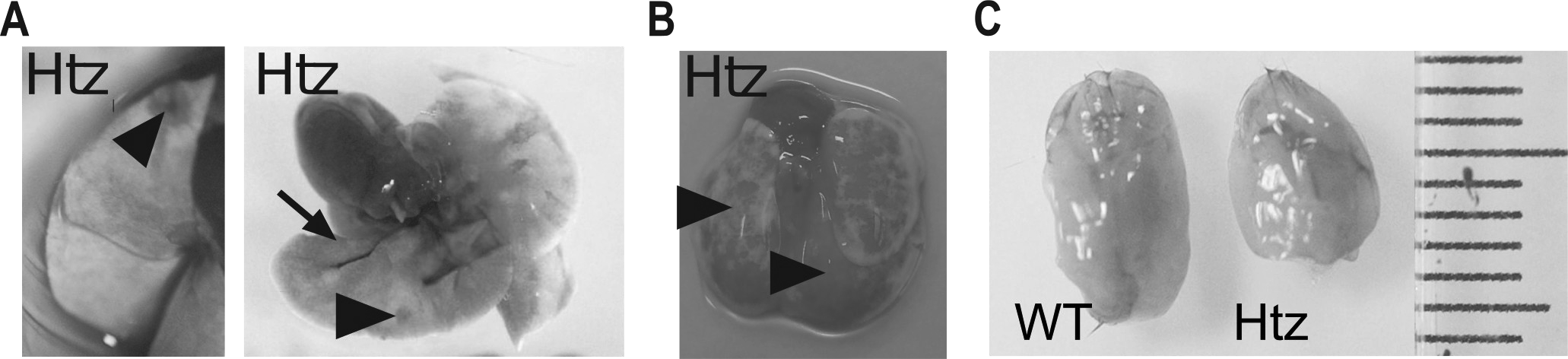
Heterozygous mice displayed abnormal lung morphology and atrophy of the salivary glands. **(A)** Macroscopic examination of a lung from a Muc5b^ko/+^ (Htz) mouse without respiratory distress. Arrowheads indicate areas of hemorrhage. (B) The hemorrhagic area is larger for a mouse with respiratory distress. (C) Macroscopic examination of salivary glands from wild-type (WT) and Htz mice. Htz mice exhibited atrophy of one salivary gland.

**Figure S6.**
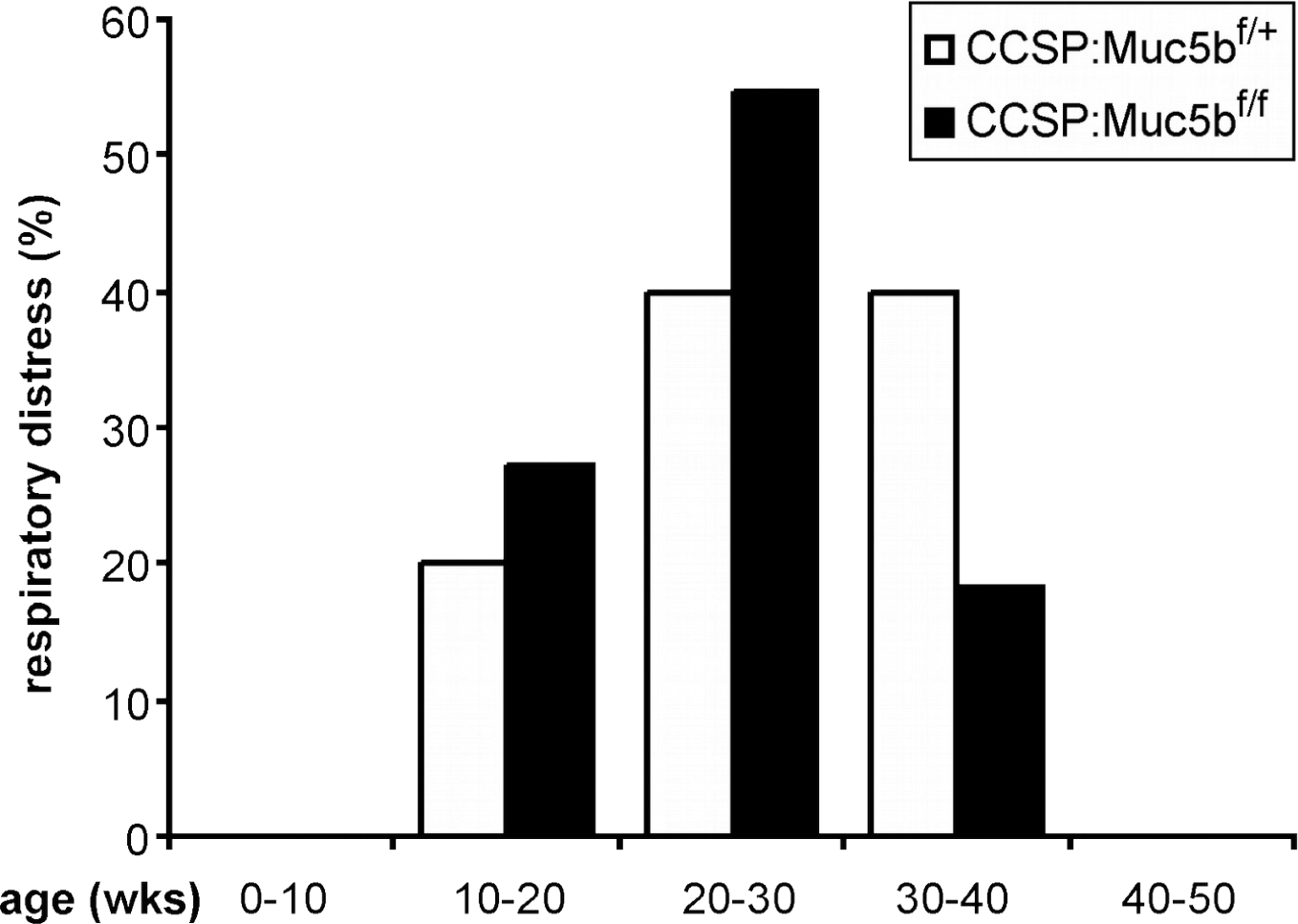
Frequency (%) of CCSPCre-positive mice with respiratory distress and carrying either one or two Muc5b-floxed alleles.

**Figure S7.**
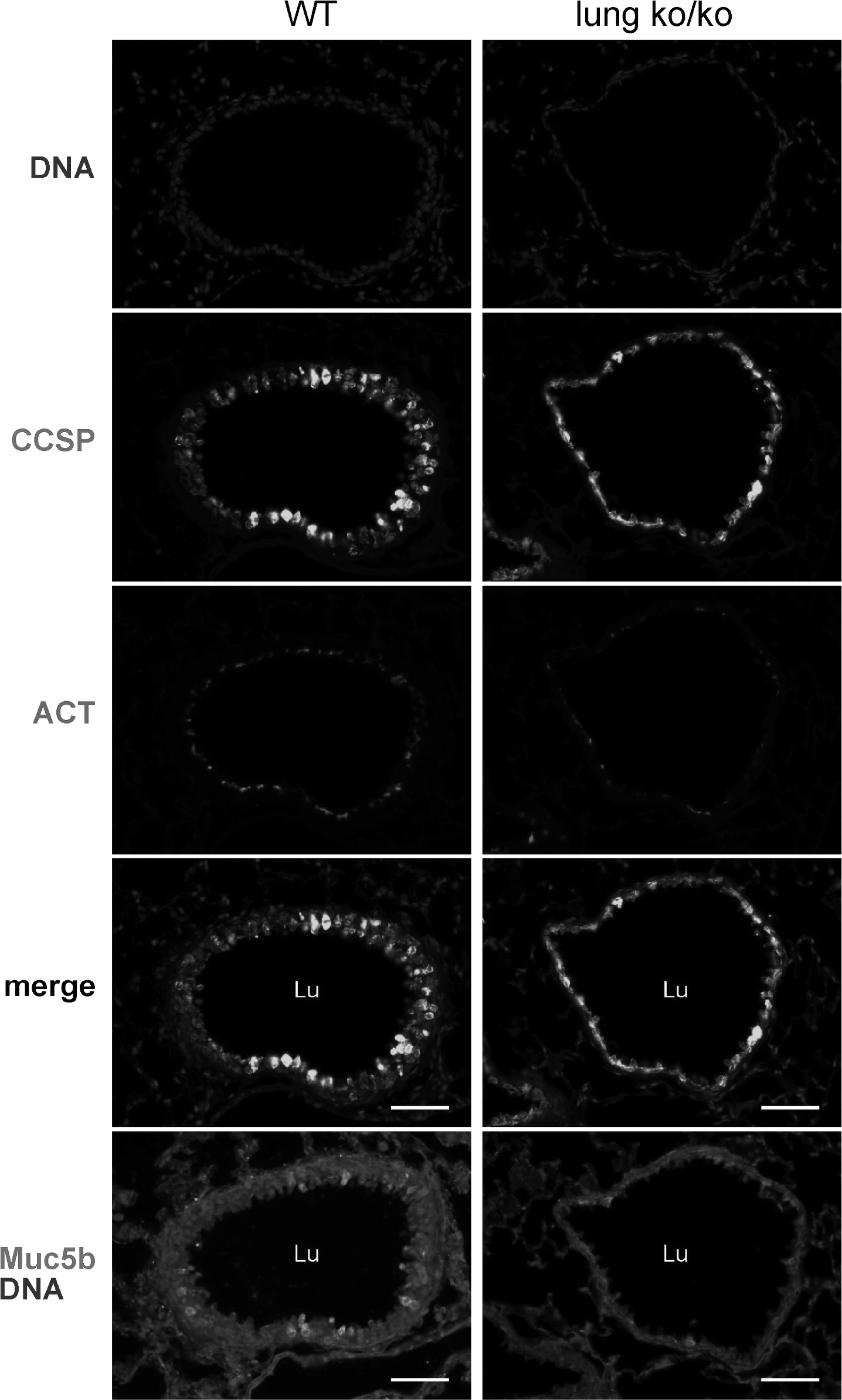
Immunohistochemical analysis of Club cells and ciliated cells. Serial bronchial sections of wild-type (WT) and Muc5b^lung ko^ mice were analyzed by immunohistochemical analysis for the detection of CCSP-, ACT- and Muc5b-positive cells. Lu = lumen; Scale bars = 50 μm.

**Video 1. Adult Muc5b^ko/+^ mouse with respiratory distress showing hunched posture, reduced locomotor activity and polypnoea.**

**Video 2. Adult Muc5b^ko/+^ mouse (different from video 1) with respiratory distress showing hunched posture, squeaking and discreet cough.**

## REFERENCES

Bertin G, Poujeol C, Rubera I, Poujeol P & Tauc M (2005). In vivo Cre/loxP mediated recombination in mouse Clara cells. Transgenic Res 14, 645–654.

Boase NA, Rychkov GY, Townley SL, Dinudom A, Candi E, Voss AK, Tsoutsman T, Semsarian C, Melino G, Koentgen F, Cook DI & Kumar S (2011). Respiratory distress and perinatal lethality in Nedd4-2-deficient mice. Nat Commun 2, 287.

Boucherat O, Chakir J & Jeannotte L (2012). The loss of Hoxa5 function promotes Notch-dependent goblet cell metaplasia in lung airways. Biol Open 1, 677–691.

Buisine MP, Desseyn JL, Porchet N, Degand P, Laine A & Aubert JP (1998). Genomic organization of the 3’- region of the human MUC5AC mucin gene: additional evidence for a common ancestral gene for the 11p15.5 mucin gene family. Biochem J 332, 729–738.

Buisine MP, Devisme L, Copin MC, Durand-Reville M, Gosselin B, Aubert JP & Porchet N (1999). Developmental mucin gene expression in the human respiratory tract. Am J Respir Cell Mol Biol 20, 209–218.

Buisine MP, Devisme L, Maunoury V, Deschodt E, Gosselin B, Copin MC, Aubert JP & Porchet N (2000). Developmental mucin gene expression in the gastroduodenal tract and accessory digestive glands. I. Stomach. A relationship to gastric carcinoma.

Camelo A, Dunmore R, Sleeman MA & Clarke DL (2014). The epithelium in idiopathic pulmonary fibrosis: breaking the barrier. Front Pharmacol 4, 173.

DeMayo FJ (1999). Advantages and pitfalls of transgenic and mutant animals. Am J Kidney Dis 33, 598–600.

Desseyn JL (2009). Mucin CYS domains are ancient and highly conserved modules that evolved in concert. Mol Phylogenet Evol 52, 284–292.

Desseyn JL, Aubert JP, Porchet N & Laine A (2000). Evolution of the large secreted gel-forming mucins. Mol Biol Evol 17, 1175–1184.

Desseyn JL, Aubert JP, Van Seuningen I, Porchet N & Laine A (1997). Genomic organization of the 3’ region of the human mucin gene MUC5B. J Biol Chem 272, 16873–16883.

Desseyn JL, Buisine MP, Porchet N, Aubert JP & Laine A (1998). Genomic organization of the human mucin gene MUC5B. cDNA and genomic sequences upstream of the large central exon. J Biol Chem 273, 30157–30164.

Desseyn JL & Laine A (2003). Characterization of mouse muc6 and evidence of conservation of the gel-forming mucin gene cluster between human and mouse. Genomics 81, 433–436.

Fahy J V & Dickey BF (2010). Airway mucus function and dysfunction. N Engl J Med 363, 2233–2247.

Fingerlin TE et al. (2013). Genome-wide association study identifies multiple susceptibility loci for pulmonary fibrosis. Nat Genet 45, 613–620.

Gustafson T, Ahlman-Hoglund A, Nilsson K, Strom K, Tornling G & Toren K (2007). Occupational exposure and severe pulmonary fibrosis. Respir Med 101, 2207–2212.

Jeffery PK (1998). Structural and inflammatory changes in COPD: a comparison with asthma. Thorax 53, 129–136.

Kamio K, Matsushita I, Hijikata M, Kobashi Y, Tanaka G, Nakata K, Ishida T, Tokunaga K, Taguchi Y, Homma S, Nakata K, Azuma A, Kudoh S & Keicho N (2005). Promoter analysis and aberrant expression of the MUC5B gene in diffuse panbronchiolitis. Am J Respir Crit Care Med 171, 949–957.

Kesimer M, Ehre C, Burns KA, Davis CW, Sheehan JK & Pickles RJ (2013). Molecular organization of the mucins and glycocalyx underlying mucus transport over mucosal surfaces of the airways. Mucosal Immunol 6, 379–392.

Li Y, Jiang D, Liang J, Meltzer EB, Gray A, Miura R, Wogensen L, Yamaguchi Y & Noble PW (2011). Severe lung fibrosis requires an invasive fibroblast phenotype regulated by hyaluronan and CD44. J Exp Med 208, 1459–1471.

Livraghi-Butrico A, Grubb BR, Wilkinson KJ, Volmer AS, Burns KA, Evans CM, O’Neal WK & Boucher RC (2017). Contribution of mucus concentration and secreted mucins Muc5ac and Muc5b to the pathogenesis of muco-obstructive lung disease. Mucosal Immunol 10, 395–407.

Madtes DK, Elston AL, Hackman RC, Dunn AR & Clark JG (1999). Transforming growth factor-alpha deficiency reduces pulmonary fibrosis in transgenic mice. Am J Respir Cell Mol Biol 20, 924–934.

Mahler CM, Berard M, Feinstein R, Gallagher A, Illgen-Wilcke B, Pritchett-Corning K & Raspa M (2014). FELASA recommendations for the health monitoring of mouse, rat, hamster, guinea pig and rabbit colonies in breeding and experimental units. Lab Anim 48, 178–192.

Noth I et al. (2013). Genetic variants associated with idiopathic pulmonary fibrosis susceptibility and mortality: a genome-wide association study. Lancet Respir Med 1, 309–317.

Phan SH (2002). The myofibroblast in pulmonary fibrosis. Chest 122, 286S–289S.

Portal C, Gouyer V, Gottrand F & Desseyn JL (2017a). Preclinical mouse model to monitor live Muc5b-producing conjunctival goblet cell density under pharmacological treatments ed. Madigan M. PLoS One 12, e0174764.

Portal C, Gouyer V, Magnien M, Plet S, Gottrand F & Desseyn JL (2017b). In vivo imaging of the Muc5b gel-forming mucin. Sci Rep 7, 44591.

Reynolds SD, Mango GW, Gelein R, Boe IM, Lund J & Stripp BR (1999). Normal function and lack of fibronectin accumulation in kidneys of Clara cell secretory protein/uteroglobin deficient mice. Am J Kidney Dis 33, 541–551.

Reynolds SD, Reynolds PR, Pryhuber GS, Finder JD & Stripp BR (2002). Secretoglobins SCGB3A1 and SCGB3A2 define secretory cell subsets in mouse and human airways. Am J Respir Crit Care Med 166, 1498–1509.

Rocco PR, Dos SC & Pelosi P (2009). Lung parenchyma remodeling in acute respiratory distress syndrome. Minerva Anestesiol 75, 730–740.

Rose MC & Voynow JA (2006). Respiratory tract mucin genes and mucin glycoproteins in health and disease. Physiol Rev 86, 245–278.

Roy MG et al. (2014). Muc5b is required for airway defence. Nature 505, 412–416.

Ryerse JS, Hoffmann JW, Mahmoud S, Nagel BA & DeMello DE (2001). Immunolocalization of CC10 in Clara cells in mouse and human lung. Histochem Cell Biol 115, 325–332.

Scotton CJ & Chambers RC (2007). Molecular targets in pulmonary fibrosis: the myofibroblast in focus. Chest 132, 1311–1321.

Seibold MA et al. (2011). A common MUC5B promoter polymorphism and pulmonary fibrosis. N Engl J Med 364, 1503–1512.

Selman M & Pardo A (2002). Idiopathic pulmonary fibrosis: an epithelial/fibroblastic cross-talk disorder. Respir Res 3, 3.

Stock CJ, Sato H, Fonseca C, Banya WA, Molyneaux PL, Adamali H, Russell AM, Denton CP, Abraham DJ, Hansell DM, Nicholson AG, Maher TM, Wells AU, Lindahl GE & Renzoni EA (2013). Mucin 5B promoter polymorphism is associated with idiopathic pulmonary fibrosis but not with development of lung fibrosis in systemic sclerosis or sarcoidosis. Thorax 68, 436–441.

Thornton DJ, Rousseau K & McGuckin MA (2008). Structure and function of the polymeric mucins in airways mucus. Annu Rev Physiol 70, 459–486.

Tompkins DH, Besnard V, Lange AW, Wert SE, Keiser AR, Smith AN, Lang R & Whitsett JA (2009). Sox2 is required for maintenance and differentiation of bronchiolar Clara, ciliated, and goblet cells. ed. Verfaillie CM. PLoS One 4, e8248.

Valque H, Gouyer V, Gottrand F & Desseyn JL (2012). MUC5B leads to aggressive behavior of breast cancer MCF7 cells. PLoS One 7, e46699.

Valque H, Gouyer V, Husson M-O, Gottrand F & Desseyn JL (2011). Abnormal expression of Muc5b in Cftr-null mice and in mammary tumors of MMTV-ras mice. Histochem Cell Biol 136, 699–708.

Wuyts WA, Agostini C, Antoniou KM, Bouros D, Chambers RC, Cottin V, Egan JJ, Lambrecht BN, Lories R, Parfrey H, Prasse A, Robalo-Cordeiro C, Verbeken E, Verschakelen JA, Wells AU & Verleden GM (2013). The pathogenesis of pulmonary fibrosis: a moving target. Eur Respir J 41, 1207–1218.

Wynn TA (2011). Integrating mechanisms of pulmonary fibrosis. J Exp Med 208, 1339–1350.

Zhang Y, Noth I, Garcia JG & Kaminski N (2011). A variant in the promoter of MUC5B and idiopathic pulmonary fibrosis. N Engl J Med 364, 1576–1577.

Zhang Z, Kundu GC, Yuan CJ, Ward JM, Lee EJ, DeMayo F, Westphal H & Mukherjee AB (1997). Severe fibronectin-deposit renal glomerular disease in mice lacking uteroglobin. Science 276, 1408–1412.

## SUPPLEMENTARY REFERENCES

Chen Y, Zhao YH & Wu R (2001). In silico cloning of mouse Muc5b gene and upregulation of its expression in mouse asthma model. Am J Respir Cell Mol Biol 164, 1059–1066.

Desseyn JL, Guyonnet-Duperat V, Porchet N, Aubert JP & Laine A (1997). Human mucin gene MUC5B, the 10.7-kb large central exon encodes various alternate subdomains resulting in a super-repeat. Structural evidence for a 11p15.5 gene family. J Biol Chem 272, 3168–3178.

Desseyn JL & Laine A (2003). Characterization of mouse mMuc6 and evidence of conservation of the gel-forming mucin gene cluster between human and mouse. Genomics 81, 433–436.

Gouyer V, Leir SH, Tetaert D, Liu Y, Gottrand F, Harris A & Desseyn JL (2010). The characterization of the first anti-mouse Muc6 antibody shows an increased expression of the mucin in pancreatic tissue of Cftr-knockout mice. Histochem Cell Biol 133, 517–525.

Howe DG, Clarke CM, Yan H, Willis BS, Schneider DA, McKnight GS & Kapur RP (2006). Inhibition of protein kinase A in murine enteric neurons causes lethal intestinal pseudo-obstruction. J Neurobiol 66, 256–272.

Leneuve P, Colnot S, Hamard G, Francis F, Niwa-Kawakita M, Giovannini M & Holzenberger M (2003). Cre-mediated germline mosaicism: a new transgenic mouse for the selective removal of residual markers from tri-lox conditional alleles. Nucleic Acids Res 31, e21.

